# Cuticle development and underlying transcriptome-metabolome associations during early seedling establishment

**DOI:** 10.1101/2022.12.09.519822

**Authors:** Keting Chen, Rupam Kumar Bhunia, Colton McNinch, Grace Campidilli, Ahmed Hassan, Ling Li, Basil J. Nikolau, Marna D. Yandeau-Nelson

**Affiliations:** Department of Genetics, Development & Cell Biology, Iowa State University, Ames, Iowa; Bioinformatics & Computational Biology Graduate Program, Iowa State University, Ames, Iowa; Roy J. Carver Department of Biochemistry, Biophysics & Molecular Biology, Iowa State University, Ames, Iowa; Molecular, Cellular, and Developmental Biology Graduate Program, Iowa State University, Ames, Iowa; Undergraduate Genetics Major, Iowa State University, Ames, Iowa; Undergraduate Data Science Major, Iowa State University, Ames, Iowa; Department of Biological Sciences, Mississippi State University, Mississippi State, Mississippi; Center for Metabolic Biology, Iowa State University, Ames, Iowa

## Abstract

The plant cuticle is a complex extracellular lipid barrier that provides protection from numerous environmental stressors and is critical for normal organ development. In this study, we investigated cuticle deposition by integrating metabolomics and transcriptomics data gathered from six different maize seedling organs of four genotypes, the inbred lines B73 and Mo17, and their reciprocal hybrids. These datasets captured the developmental transition of the seedling from heterotrophic skotomorphogenic growth to autotrophic photomorphogenic growth, which is a transition that is highly vulnerable to environmental stresses. Statistical interrogation of these data reveals that the predominant determinant of cuticle composition is seedling organ type, whereas the seedling genotype has a smaller effect on this phenotype. Gene-to-metabolite associations assessed by joint statistical analyses of transcriptome and metabolome datasets identified three gene networks connected with the deposition of different fractions of the cuticle: a) cuticular waxes; b) cutin of aerial organs and suberin of roots; and c) both of these fractions. These networks consist of genes that encode known components of the machinery that supports cuticle deposition, demonstrating the utility of this integrated omics approach. Moreover, these gene networks reveal three additional metabolic programs that appear to support cuticle deposition, including processes of a) chloroplast biogenesis, b) lipid metabolism, and c) molecular regulation (e.g., transcription factors, post-translational regulators and phytohormones). This study demonstrates the wider physiological metabolic context that can determine cuticle deposition and lays the groundwork for new targets for modulating properties of this protective barrier.

## INTRODUCTION

One of the key evolutionary developments that enabled plants to colonize the terrestrial environment appears to have been the ability to synthesize an external hydrophobic barrier, the cuticle (Seale, 2020). Although the primary functionality of the cuticle is as a water-barrier, additional attributes assigned to the cuticle include protection from many biotic (e.g., bacterial, fungal and insect pests) and abiotic (e.g., drought and UV-radiation) environmental stressors (Shepherd and Wynne Griffiths, 2006; Yeats and Rose, 2013).

On aerial organs of terrestrial plants, the cuticle is composed of a cutin matrix, which is immersed with and coated by solvent-extractable cuticular waxes. The cutin matrix is primarily a polyester polymer comprising epoxy- and hydroxy-fatty acids and glycerol, and cuticular waxes consisting of different mixtures of lipid classes, including very-long-chain fatty acids (VLCFAs), fatty aldehydes, primary and secondary fatty alcohols, wax esters, hydrocarbons, ketones and terpenes (Reynoud et al., 2021). In contrast, roots have a physiologically different function, being the organ that collects water and water-soluble nutrients from the soil matrix, thus the structure and role of cuticle-like metabolites is different from the aerial organs (Baxter et al., 2009). Specifically, a cuticle is associated with the cap of the developing roots, and a cutin-homolog (i.e., suberin) is a lipid polymer associated with the periderm and endoderm of roots that appears to function as a water-barrier, possibly facilitating water-movement (Berhin et al., 2019).

In aerial organs, cuticle biosynthesis occurs in epidermal cells, and cuticular components are deposited on the extracellular, exterior facing surfaces of the cells toward the aerial environment (Bhanot et al., 2021). Cuticle biosynthesis is developmentally regulated as manifest by different cuticle compositions among different organs of the plant, or even different tissues within the same organ. For example, in maize, fatty alcohols constitute >60% of the cuticular waxes on leaves of juvenile identity, but they account for only ∼15% on adult-identity leaves (Avato et al., 1987). These compositional differences are juxtaposed with increased abundances of wax esters (from ∼15% to 40%) and hydrocarbons (from 1% to ∼20%). More recent studies have revealed that on developing adult-identity maize leaves, the predominant wax components differ according to position on the leaf, with hydrocarbons predominant at the leaf base and alkyl esters at the leaf tip (Bourgault et al., 2020). In contrast, the stigmatic silks of maize express a very different composition of cuticular waxes, being rich in both saturated and unsaturated hydrocarbons (often exceeding 90% of the cuticular waxes), with minor amounts of VLCFAs, fatty aldehydes and fatty alcohols (Perera et al., 2010; Loneman et al., 2017; Dennison et al., 2019).

Both the cutin and cuticular wax biosynthetic pathways utilize 16- or 18-carbon chain-length fatty acyl-CoAs as the initial precursors. Cutin monomers are generated via hydroxylation of these fatty acyl-CoAs, which can occur at multiple positions on the alkyl-chain (i.e., at the 2-position, midchain or at the ω-position) and some of these hydroxy-fatty acids can be converted to epoxy-fatty acids (Fich et al., 2016). The hydroxylated-FAs are acylated to glycerol-3-phosphate to form the cutin polyester precursors (Li et al., 2007), which are transported extracellularly for the final cutin polymerization reaction catalyzed by cutin synthase (Yeats and Rose, 2013; Fich et al., 2016).

The cuticular wax components are generated from very-long chain fatty acyl-CoAs, which are products of the endoplasmic reticulum (ER)-bound fatty acid elongase (FAE) system that iteratively adds 2-carbon units per cycle to preexisting C16 or C18 fatty acyl-CoAs (Yeats and Rose, 2013; Campbell et al., 2019). Downstream of FAE, two parallel pathways generate primary fatty alcohols, fatty aldehydes and wax esters (i.e., the reductive pathway), or primary aldehydes, hydrocarbons, secondary fatty alcohols and ketones (i.e., the decarbonylative pathway). Notably, these two pathways can utilize very long chain fatty acyl-CoAs of a variety of chain lengths (20-34 carbons and possibly longer), generating a homologous series of VLCFA-derivatives as a mixture of cuticular wax metabolites (Kolattukudy and Walton, 1973; Yeats and Rose, 2013).

In this study, transcriptome profiles coupled with cutin/suberin monomer and cuticular wax profiles were gathered from six seedling organs of four maize genotypes (the inbred lines B73 and Mo17, and the reciprocal hybrids B73×Mo17 and Mo17×B73) representing a spatial gradient that captures the transition of plant growth mode from heterotrophic skotomorphogenesis (i.e., seedling development in darkness) to autotrophic photomorphogenesis. The gene networks underlying the changes in cuticle composition during this transition were queried by a multi-omics integration approach designed to assess the joint statistical association between transcriptomes and metabolomes. This approach identified genetic networks comprising ∼1,900 genes associated with cutin/suberin monomer and/or cuticular wax deposition. These gene networks were composed of not only genes known to be components of cuticle biosynthetic pathways, but also identified many genes, and thereby biological processes, not previously recognized as being associated with cuticle deposition, including post-translational regulation, chloroplast development, lipid degradation pathways and signaling mechanisms.

## RESULTS

### Cuticular wax composition of seedling organs from different genetic backgrounds

Cuticular waxes were profiled from six seedling organs (i.e., roots, coleoptiles, first leaf sheathes, leaves encased by the first leaf sheath (encased leaves), and the first and second leaf blades) (Fig. 1A) of four different maize genotypes (i.e., the two inbreds, B73 and Mo17, and the reciprocal hybrids, B73×Mo17 and Mo17×B73). These analyses identified six classes of cuticular wax components: VLCFAs, fatty alcohols, fatty aldehydes, wax esters and hydrocarbons, and a small quantity of terpenes, the latter being particularly abundant in roots (Fig. 1B). The alkyl cuticular wax components ranged between 12- and 34-carbon atoms in chain length, with the predominant chain length occurring at 32-carbons for fatty acids, fatty alcohols and fatty aldehydes and 31-carbons for alkanes and alkenes (Supplemental Fig. S1A and Table S1A). The range of chain lengths, however, were slightly different among the different classes of cuticular wax components (e.g., fatty alcohols ranged between 18 and 34 carbon atoms, whereas the fatty aldehydes ranged between 26 and 34 carbon atoms) (Supplemental Fig. S1A and Table S1A).

**Figure 1.**
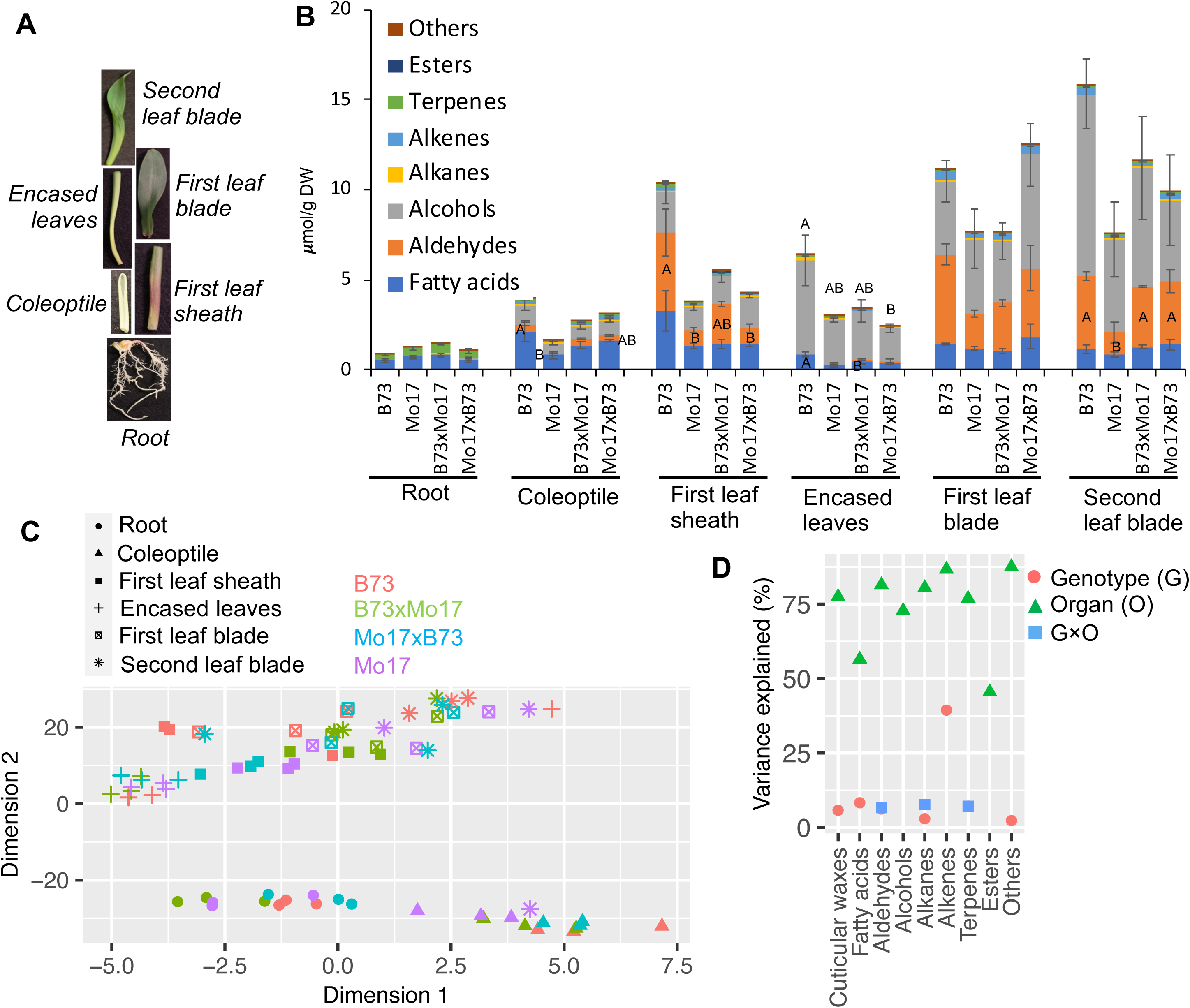
Comparison of cuticular waxes among seedling organs from maize inbreds and reciprocal hybrids. (A) Seedling organs collected for metabolomics and transcriptomics analysis. (B) The accumulation of cuticular wax lipid classes among the organs of different genotypes. Statistically significant differences (calculated by Tukey’s HSD test; p-value <0.05) in the accumulation of different cuticular wax lipid classes among different genotypes for a given organ are indicated by different letters; (C) tSNE visualization of different cuticular wax metabolomes expressed by triplicate analysis of different organs from different genotypes (D) The impact of genotype, organ, and genotype x organ interactions on different cuticular wax lipid classes calculated as partial R^2^ values by Type I ANOVA. Factors that do not significantly impact variation on the cuticular wax lipid classes (p-values <0.05) are absent from the figure.

Regardless of the genotype that was evaluated, first and second leaf blades accumulated the largest amounts of cuticular waxes, whereas the first leaf sheath and the encased leaves accumulated approximately half that amount and minimal accumulation of cuticular wax-like metabolites were observed on the roots. Furthermore, the majority of the cuticular waxes that were recovered from these leaves were fatty alcohols and fatty aldehydes, which together accounted for ∼80% of the recovered cuticular wax components (Fig. 1B). The C32 fatty aldehyde is the predominant cuticular wax constituent identified on first and second leaf blades, while it is only a minor constituent on coleoptiles, first leaf sheath, encased leaves, and roots (Table S1A). In addition, cuticular wax compositions of roots and coleoptiles vary substantially from the other organs, comprising wax metabolites of shorter chain lengths than for the other organs (i.e. chain lengths of 30 carbons or shorter; Fig. S1A). Cuticular wax-like metabolites accumulate to much lower levels on the roots, and comprise approximately equal amounts of terpenes and VLCFAs. In contrast, cuticular wax metabolites accumulate to higher levels on coleoptiles and consist of primarily VLCFAs and fatty alcohols (Fig. 1B and 1C).

The relative contributions of different genotypes and organ types to the variation in cuticular wax composition was visualized via two unsupervised statistical methods, principal component analysis (PCA) and *t*-Distributed Stochastic Neighbor Embedding (tSNE) (Fig. 1C and Fig. S2A). Both visualization methods indicate that the differences in cuticular wax metabolomes are primarily determined by the different gene expression programs of the seedling organs, with minor contributions by the different gene expression programs among the four genotypes (Fig. 1C). Specifically, PC1 and PC2 that together explained 55% of the observed variance in cuticular wax composition, separated the root and coleoptile samples from the other seedling organs. Examination of each individual class of metabolites via Analysis of Variance (ANOVA) revealed that the organ type explains a large proportion of the observed variance for most cuticular wax classes (ranging between 45% and 88%), and genotype and genotype × organ interactions contribute to a far lesser extent (<10%) (Fig 1D). Collectively, these analyses demonstrate that the major influence on the cuticular wax metabolome is the seedling organ type rather than the genetic background of the seedling.

### Cutin and suberin monomer composition of seedling organs from different genetic backgrounds

In parallel to the analyses of the cuticular waxes, the six dissected organs among the inbreds and hybrids were also analyzed to determine the monomer compositions of the cuticle-associated polymeric lipids (i.e., cutin of the aerial organs and suberin of the roots). Chemical depolymerization of these polymeric lipids identified five major monomer subclasses: fatty acids, hydroxy fatty acids (i.e., with the hydroxyl groups being situated at the 2-position, the ω-position or at an unknown position) and phenolics (Fig. 2A). The fatty acids that were recovered ranged between 16 and 30 carbon atom chain lengths, whereas the ω-hydroxy fatty acids ranged between 16 and 22 carbons, and 2-hydroxy fatty acids ranged between 16 and 26 carbons (Supplemental Fig. S1B and Table S1B). The phenolic compounds that were recovered include caffeic acid, coumaric acid and ferulic acid.

**Figure 2.**
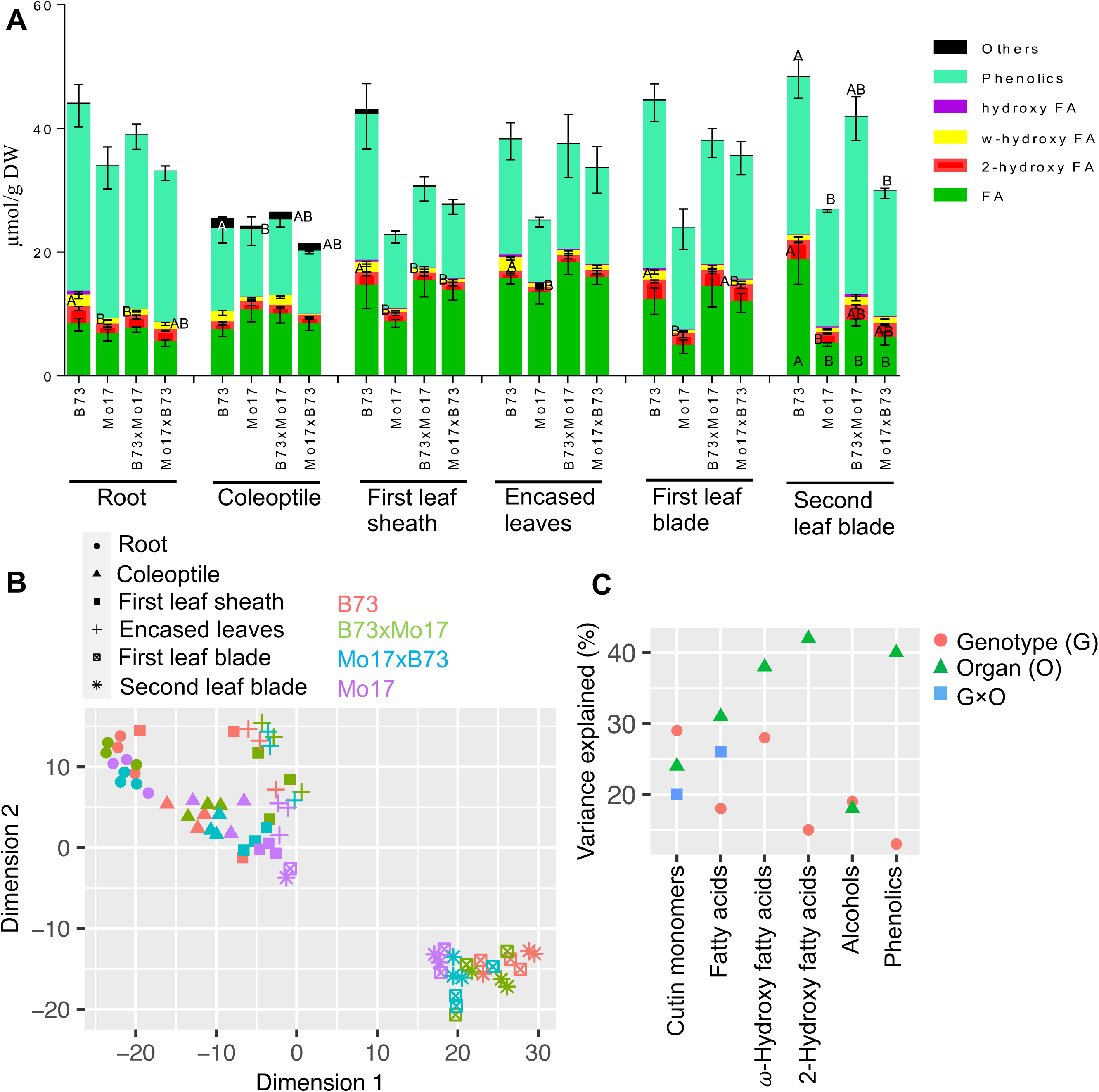
Comparison of cutin and suberin monomers among seedling organs from maize inbreds and reciprocal hybrids. (A) The accumulation of cutin and suberin monomers among the organs of different genotypes. Statistically significant differences (calculated by Tukey’s HSD test; p-value <0.05) in accumulation of monomer classes among different genotypes for a given organ are indicated by different letters; (B) tSNE visualization of cutin and suberin monomer metabolomes expressed by triplicate analysis of different organs from different genotypes; (C) Partial R^2^ values representing the impact of genotype, organ, and genotype x organ interaction on different cutin monomer classes as evaluated by Type I ANOVA. Factors without significant on the cutin/suberin monomers (p-values <0.05) are not presented in the figure.

PCA and tSNE analyses were used to visualize the effects of different genotypes and organ types on the variance of the cuticular lipid monomer compositions (Fig. 2B and Fig. S2B). In combination, PC1 and PC2, which together explained 49% of the observed variance in monomer composition, separated the root and coleoptile samples from the other seedling organs (Fig. S2B), and the higher-resolution tSNE visualization generated a more distinct separation of the two leaf blades relative to the other seedling organs (Fig. 2B). ANOVA supports the PCA and tSNE data visualizations; namely different seedling organs explain between 18% and 42% of the observed variance among the cutin/suberin monomer classes (Fig. 2C), while the genotype difference among the samples has a smaller, but still substantial impact, explaining between 13% and 29% of the observed variance in each of the metabolite classes (Fig. 2C). Finally, the interaction between genotype and organ type explains 26% of observed variance for total non-hydroxy-fatty acids (Fig. 2C). Collectively, these analyses demonstrate that analogous to the cuticular wax metabolite profiles, organ type predominantly influences cutin/suberin monomer composition, while genotype difference has a smaller effect on these profiles.

### Variation in global gene expression among seedling organs of different genotypes

The transcriptome of each seedling organ was sequenced, and the resulting transcripts were mapped to the B73 reference genome (Jiao et al., 2017), incorporating the corresponding gene sequence variants from Mo17 (Sun et al., 2018). These assemblies were used to quantify expression of 30,931 genes that are detected among the six seedling organs from the four genotypes that were evaluated. Visualization of the datasets by tSNE and PCA demonstrate that the root transcriptomes segregate into a discrete cluster, whereas the transcriptomes of the other organs are grouped into three distinct clusters, one containing the coleoptile and first leaf sheath, another containing the first and second leaf blades, and the third cluster encompasses the encased leaves (Fig. S3). However, the transcriptomes within these clusters exhibit no consistent separation based on genotype.

Differentially expressed gene (DEG) analysis of the data was performed to compare the transcriptomes between every pair of genotypes (i.e., six comparisons) for each organ. The total non-redundant DEGs between every pair of genotypes (Fig. S4) comprises <8% of the entire transcriptome, with the majority of the DEGs identified in each organ occurring between the inbreds B73 and Mo17, and between the combined hybrids and one of the parental inbreds, while the hybrids B73×Mo17 and Mo17×B73 express near identical transcriptomes, with only one detectable DEG between them (Fig. S4). In contrast, the total non-redundant DEGs between every pair of organs (i.e., 15 comparisons) comprises 34% to 46% of the entire transcriptome in each genotype (Fig. S5). Collectively, these data demonstrate that the gene expression program is primarily driven by the developmental program that generates the six different organs, and that these programs are highly similar among the different genotypes that were assessed.

### The association between global gene expression profiles and cuticle deposition

In the context of different genotypes and different seedling organs, two independent approaches were used to interrogate the associations between the transcriptome and cuticle composition. These analyses utilized all the gathered metabolite abundance data and 22,462 transcript abundances whose expression were at detectable levels in more than half of the samples that were sequenced. These analyses are schematically illustrated in Fig. 3, and comprised: (1) a Weighted Gene Co-expression Network Analysis – Random Forest (WGCNA-RF) approach that assessed the relationship between co-expressed gene clusters and the changes in cuticle composition, which identified gene clusters with significant collective association to cuticle composition, and (2) a multi-omics integration approach that conducted a joint statistical association analysis between individual transcripts and individual cuticle metabolites, which identified individual genes associated with differences in cuticle composition. Subsequently, the results from the two approaches were integrated to define gene networks likely underlying the observed changes in cuticle composition amongst seedling organs and between genotypes (Fig. 3).

**Figure 3.**
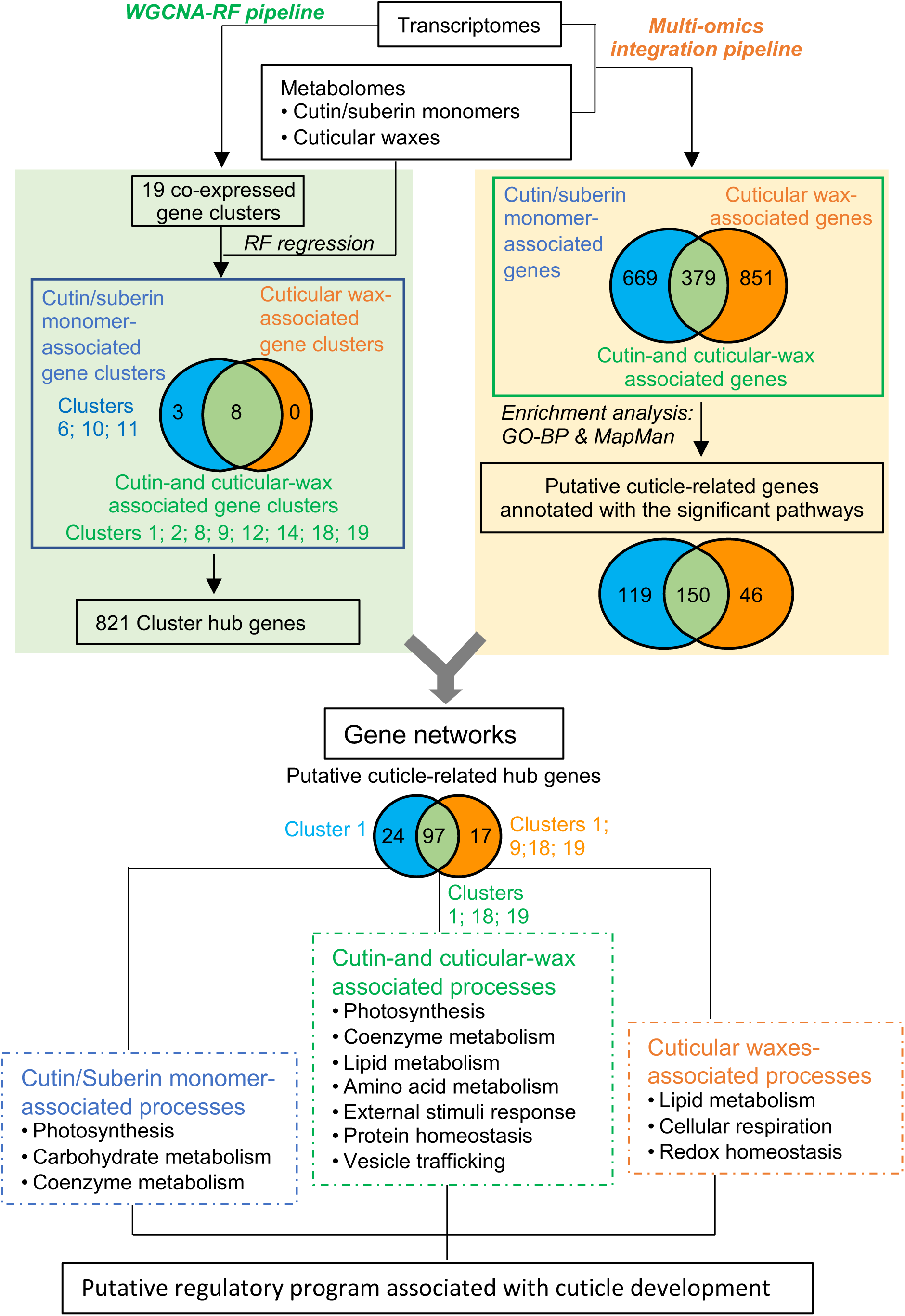
Schematic summary of the outcomes from the parallel analysis of the combined transcriptome-metabolome dataset by two independent pipelines. The WGCNA-RF pipeline identified 19 co-expression gene clusters, 3 of which are associated with the accumulation of cutin/suberin monomers and 8 are associated with the accumulation of both the cutin/suberin monomers and cuticular waxes. The multi-omics integration pipeline identified 119, 46 and 150 46 genes that are associated with cutin/suberin monomers, cuticular waxes and the combination of cutin/suberin monomers and cuticular waxes, respectively. Integrating these two outcomes generated gene networks that identify different metabolic pathways that associated with different processes that assemble the integrated cuticle.

#### Identification of co-expressed gene clusters associated with cutin/suberin monomer and cuticular wax composition by the WGCNA-RF approach

WGCNA was applied to the transcriptome datasets to first reveal clusters of genes that showed co-expression patterns among the six seedling organs from the four genotypes. Subsequently, the application of a random forest regression model revealed the association of each co-expressed gene cluster with changes in the composition of the cuticle metabolome. One can often infer that genes within a co-expression cluster that show high correlation to a trait (in this case the cuticle composition trait) have a significant role in the biological processes that determine that trait (Carlson et al., 2006).

WGCNA determined 19 co-expression gene clusters, leaving 378 genes that showed expression patterns that were not clusterable (Fig. 4 and Supplemental Table S2A). Subsequently, the expression of an eigengene was calculated for each co-expression gene cluster, representing the average gene expression profile per genotype and thereby visualizing the cluster average expression pattern among the six organs. Hierarchical clustering further categorized the 19 co-expression clusters into nine classes (Class I to IX; Supplemental Fig. S6 and Fig. 4). Genes in six of these classes (I, IV, V, VI, VII, and IX) exhibit an expression pattern that varies primarily among seedling organs, with little if any difference among the four genotypes that were evaluated. Genes within Class I clusters show highest expression levels in the aerial organs of the seedling, particularly in the leaf blades, while Class IX genes show the opposite pattern, with maximum expression occurring in roots, and minimal expression in the aerial organs (Fig. 4). Genes in Classes IV, V, and VI, show maximal gene expression in coleoptiles, first leaf sheath, or encased leaves, while Class VII genes demonstrate minimal expression in these organs (Fig. 4).

**Figure 4.**
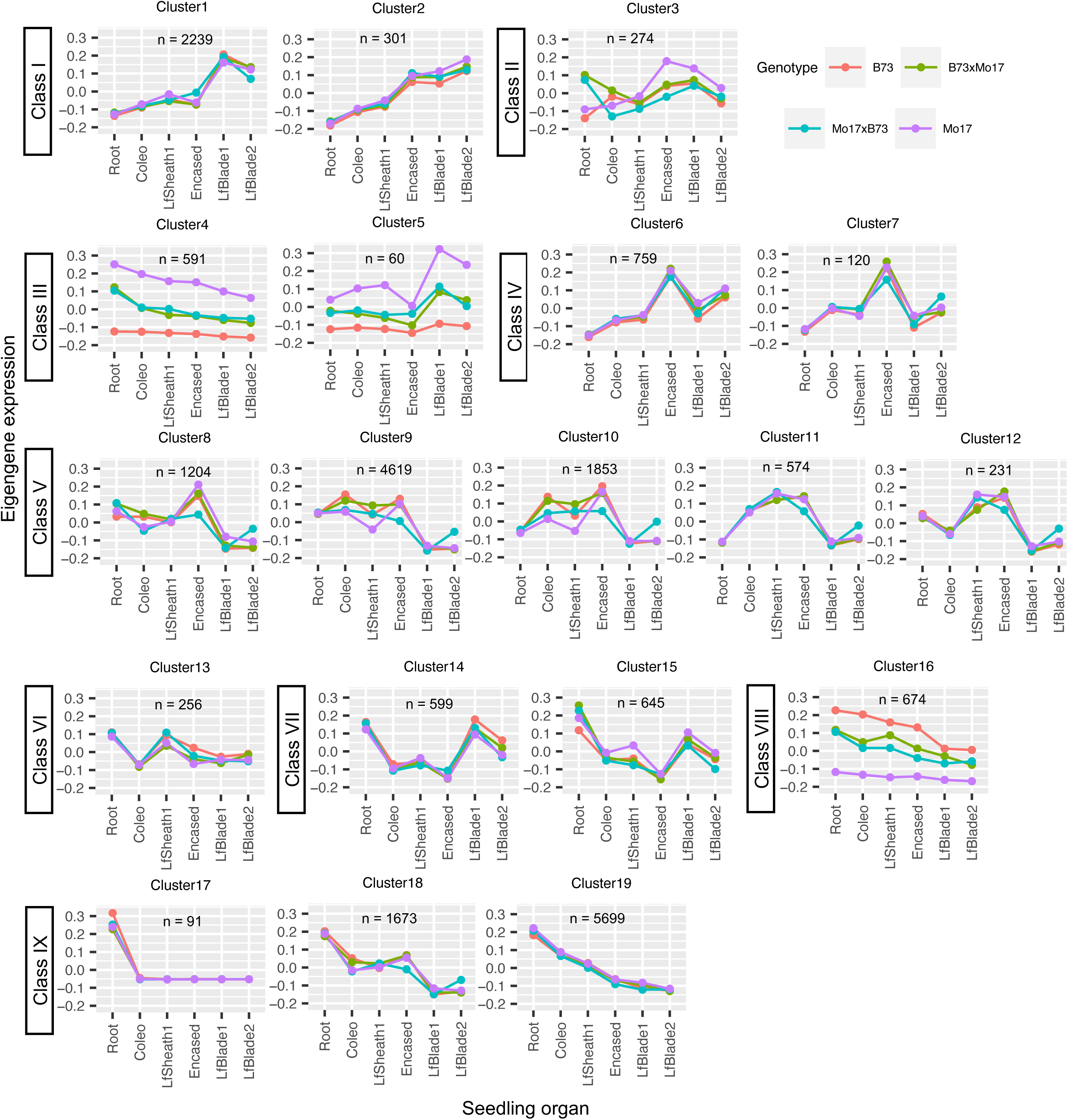
Co-expression gene clusters determined by WGCNA. Each color identifies the eigengene expression pattern that represents the co-expressed gene cluster for each maize genotype (i.e., B73, Mo17, B73×Mo17 and Mo17×B73). Hierarchical clustering based on the eigengene expression pattern categorized the 19 co-expression gene clusters into 9 classes (Class I to Class IX). Coleo, coleoptile; LfSheath1, first leaf sheath; Encased, encased leaves; LfBlade1, first leaf blade; LfBlade2, second leaf blade; n = number of genes in each co-expression gene cluster.

In contrast, genes within the Class II, III, and VIII clusters exhibit expression patterns that differ among the organs and are also affected by the genotype of these organs. Collectively these three classes contain only 1,599 genes (∼7% of the genes analyzed by WGCNA), but the developmental programs that underlie the expression patterns of these genes in each organ are differentially affected by the genotype of the seedlings. Opposite effects of genotype on Class III and Class VIII genes are observed, with the expression in B73 being lower than in Mo17 for Class III genes, whereas expression of Class VIII genes is higher in B73; and in both cases the expression in the reciprocal hybrids is indistinguishable from each other and is at the mid-point between the two parental inbreds (Fig. 4). Finally, expression of genes in Class II demonstrates a complex interaction between genotype and seedling organs; this is best illustrated by comparing the expression patterns in roots to the patterns observed in the aerial organs. In roots, higher expression is observed in the two hybrids as compared to the parental inbreds, but in the aerial organs the higher expression is observed in the Mo17 parental inbred, and expression is at a similar, lower level among the other three genotypes (Fig. 4).

The associations between specific gene co-expression clusters, and differences in cutin/suberin monomer and cuticular wax compositions were interrogated by a random forest regression model. This model used the calculated expression of the eigengenes that represent each co-expression gene cluster, as well as the experimentally measured expression of the individual 378 genes that were not clusterable, to predict the previously calculated tSNE scores for cutin/suberin monomers and cuticular waxes (Figs. 1C and 2B), thereby identifying gene co-expression clusters that are associated with different cuticle compositions. The importance of each co-expression cluster to the model was evaluated by the random forest model comparisons described in the Methods. These analyses identified 11 co-expression gene clusters as significantly associated with differences in cuticle composition (Clusters 1, 2, 6, 8, 9, 10, 11, 12, 14, 18 and 19). Eight of these clusters (Clusters 1, 2, 8, 9, 12, 14, 18 and 19) were significantly associated with differences in both cuticular wax composition and cutin/suberin monomer composition, and three clusters (Clusters 6, 10, and 11) were uniquely associated with differences in cutin/suberin monomer composition; no clusters were uniquely associated with differences in cuticular wax composition (Fig. 3, Table 1, and Supplemental Table S2B and S2C). Next, we used three criteria to find “hub genes” that were representative of these 11 co-expression gene clusters associated with differences in cuticle composition: (1) high connectivity to the other genes in the cluster (i.e., gene-to-gene associations); (2) high association with the corresponding eigengene, and (3) high association with the compositional differences in cutin/suberin monomer and/or cuticular wax traits. A total of 812 hub genes were thus identified that we hypothesize play important roles in determining cuticle composition (Table 1 and Supplemental Table S2D).

**Table 1.** Co-expressed gene clusters associated with cuticle composition.

| Cluster | Number of genes | Associated cuticle fraction <sup>a</sup> | Number of putative cuticle-related genes | Gene-to-tSNE correlation threshold <sup>c</sup> | Number of hub genes | Number of hub genes that are putative cuticle-related genes |
| --- | --- | --- | --- | --- | --- | --- |
| *1*<sup>b</sup> | 2239 | Cutin/suberin;<br>Cuticular wax | 437<sup>b</sup> | 0.75 | 140 | 131 |
| 2 | 301 | Cutin/suberin;<br>Cuticular wax | 26 | 0.7 | 15 | 10 |
| 6 | 759 | Cutin/suberin | 12 | 0.6 | 35 | 1 |
| 8 | 1204 | Cutin/suberin;<br>Cuticular wax | 6 | 0.5 | 58 | 0 |
| 9 | 4619 | Cutin/suberin;<br>Cuticular wax | 139 | 0.65 | 131 | 41 |
| 10 | 1853 | Cutin/suberin | 13 | 0.55 | 122 | 0 |
| 11 | 574 | Cutin/suberin | 17 | 0.45 | 15 | 1 |
| 12 | 231 | Cutin/suberin;<br>Cuticular wax | 1 | 0.3 | 7 | 0 |
| 14 | 599 | Cutin/suberin;<br>Cuticular wax | 3 | 0.3 | 5 | 0 |
| 18 | 1673 | Cutin/suberin;<br>Cuticular wax | 107 | 0.65 | 108 | 52 |
| *19*<sup>b</sup> | 5699 | Cutin/suberin;<br>Cuticular wax | 1003<sup>b</sup> | 0.75 | 185 | 179 |

<sup>a</sup> WGCNA-based co-expression clusters are listed as being significantly associated with either or both cutin/suberin monomer and cuticular wax compositions. Evaluation of the association between gene clusters and metabolome compositions is detailed in Methods.
<sup>b</sup> Clusters that are significantly enriched (corrected p-value <0.01 according to Fisher's exact test) with the putative cuticle-related genes selected by the multi-omics integration analysis.
<sup>c</sup> Different threshold for the correlations between individual gene expression and tSNE scores of cutin/suberin monomers and/or cuticular waxes are selected for each module such that ~5% of genes are identified as hub genes.

#### Identification of putative cuticle-related genes associated with cutin/suberin monomer and cuticular wax composition by multi-omics integration

Synergistic to the WGCNA-RF approach that identified hub genes, a complementary series of gene-metabolite association approaches were implemented to identify genes that may mediate cuticle formation. Specifically, three multivariate statistical models were employed for joint statistical analysis of the metabolome and transcriptome datasets, which aimed to identify individual genes whose expression patterns are associated with the accumulation patterns of cutin/suberin monomers and cuticular wax metabolites. These statistical models included partial least square regression (PLS), sparse partial least square regression (sPLS) and random generalized linear model (rGLM). PLS identified genes that were highly expressed (i.e., >100 FPKM), whereas sPLS and rGLM identified genes with lower expression levels (i.e., >0 and <100 FPKM). In the PLS and sPLS models, the concentrations of all individual cutin/suberin monomer or cuticular wax metabolites were used as the response variables, whereas in the rGLM, the tSNE components of these metabolites were used as the response variables (Fig. 1C and Fig. 2B). The resultant list of putative cuticle-related genes (i.e., the explanatory variables) comprised ∼1,900 genes (listed in Supplemental Table S3) that were selected by any single statistical model. This gene list includes 669 genes that are uniquely associated with cutin/suberin monomers (cutin/suberin monomer-associated genes), and 78 of these had been identified as hub genes by the WGCNA-RF approach (Supplemental Table S3). Additionally, 851 genes are uniquely associated with cuticular wax metabolites (cuticular wax-associated genes), which includes 152 hub genes identified by the WGCNA-RF approach. Finally, 379 genes have associations with both cutin/suberin monomers and cuticular waxes, and this list includes 185 hub genes identified by the WGCNA-RF approach (Fig. 3).

Each of these putative cuticle-related gene lists were parsed into functional categories by performing two types of enrichment analyses: 1) gene ontology enrichment of biological processes (GO-BP terms) (Ashburner et al., 2000), that were further trimmed using REVIGO (Supek et al., 2011); and 2) enrichment into MapMan function bins (Schwacke et al., 2019). The former descriptors identify the participation of individual genes in broad categories of biological processes, whereas the latter assign individual genes to different metabolic pathways.

Cross-referencing the results of MapMan analysis (Supplemental Table S4) and GO-BP analysis (Supplemental Table S5, S6 and S7) identified ten distinct metabolic pathways that are significantly enriched among the ∼1,900 putative cuticle-related genes. Specifically, 119 of the 699 cutin/suberin monomer-associated genes (Supplemental Table S4 and S5) are significantly enriched with three pathways, “Photosynthesis”, “Carbohydrate metabolism”, and “Coenzyme metabolism” (Fig. 3). Parallel analyses of the 852 cuticular wax-associated genes identified a subset of 46 genes that were significantly enriched with three pathways, “Cellular respiration”, “Lipid metabolism”, and “Redox homeostasis” (Fig. 3, Supplemental Table S4 and S6). Finally, 150 of the 379 genes that are both cutin- and cuticular wax-associated are significantly enriched with seven pathways, “Amino acid metabolism”, “Coenzyme metabolism”, “External stimuli response”, “Lipid metabolism”, “Photosynthesis”, “Protein homeostasis”, and “Vesicle trafficking” (Fig. 3, Supplemental Table S4 and S7).

#### Deduction of co-expression networks from the combination of hub genes and putative cuticle-related genes

Co-expression gene networks were constructed by integrating the co-expressed clusters obtained from the WGCNA-RF approach that identified 821 hub genes, with the list of 315 putative cuticle-related genes identified by the multi-omics integration approach and refined by gene function enrichment analyses (Supplemental Table S4, S5, S6, and S7) (Fig. 3). This integration enabled the deduction of three gene networks, two of them being specific for either cutin/suberin monomers or cuticular waxes, respectively, and a third gene network that underlies both cutin/suberin monomers and cuticular waxes (Fig. 5).

**Figure 5.**
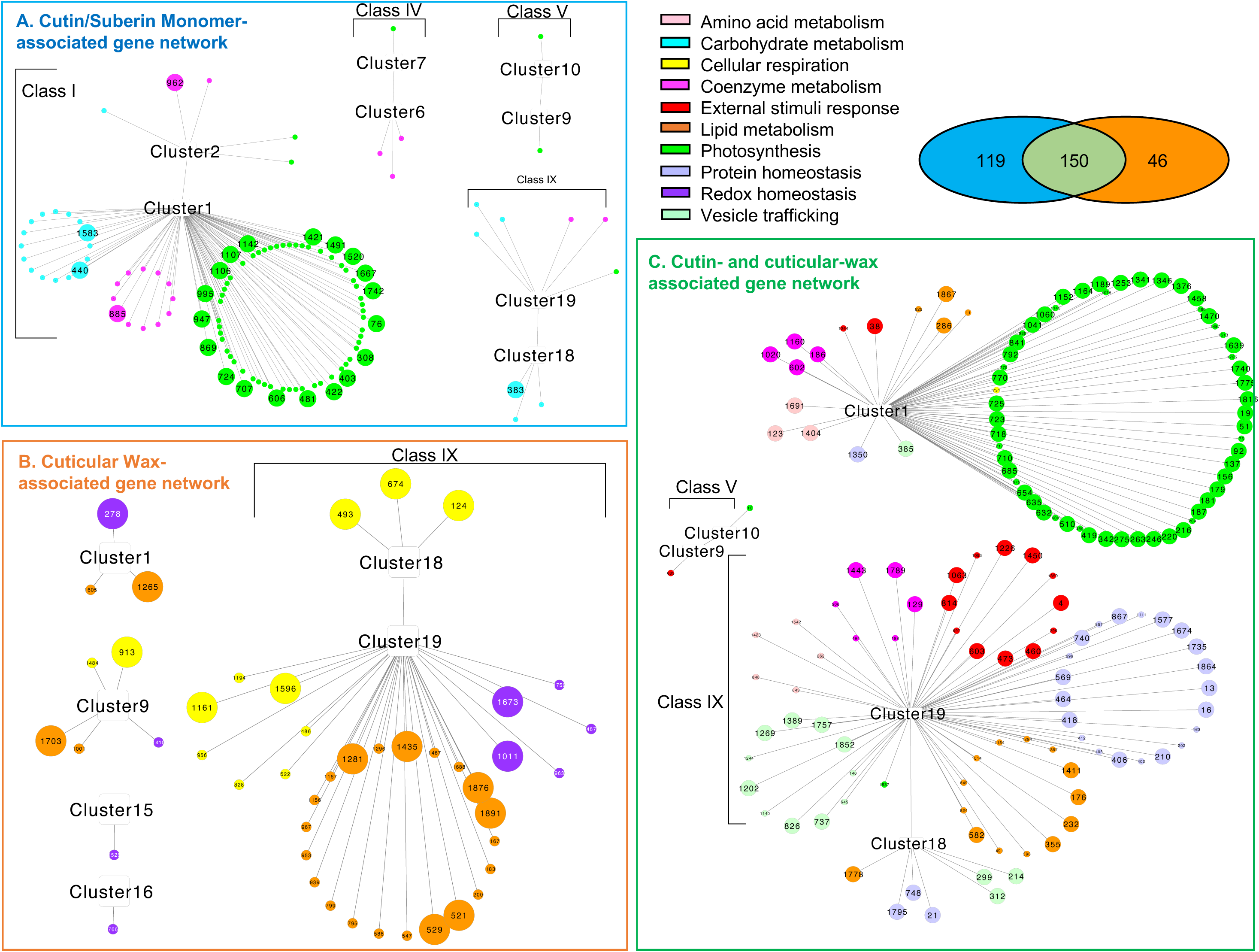
Co-expression gene networks identified by the combined WGCNA-RF and multi-omics integration pipelines. The three gene networks incorporate: a) the cutin/suberin monomer-associated genes; b) cuticular wax associated genes; and c) cutin/suberin and cuticular waxes associated genes. Numerically labelled colored symbols represent genes, whose identity is listed in Supplemental Table S4; colors of these symbols identify different MapMan metabolic function bins. Larger sized symbols represent the hub genes identified by the WGCNA-RF pipeline.

The cutin/suberin monomer-associated gene network identified by the multi-omics approach consists of 119 genes belonging to eight co-expression clusters that were identified by the WGCNA approach (Fig. 5). Most of the genes in this network (∼87%) reside in Class I co-expression clusters (i.e., Clusters 1 and 2 of Fig. 4), wherein gene expression is highest in the leaf blades of the seedlings. The other genes in this network are distributed among Class IV, Class V and Class IX co-expression clusters (Fig. 4 and 5), with genes from Classes IV and V exhibiting highest expression in the aerial organs as compared to the roots. Additionally, 24 of the 119 cutin/suberin monomer-specific genes are also identified as hub genes within their resident co-expression clusters (Fig. 3 and Fig. 5). The 119 genes associated with the cutin/suberin monomer gene network were mapped to three metabolic pathways: “Photosynthesis”, “Coenzyme metabolism”, and “Carbohydrate metabolism”.

Similar analyses identified a cuticular wax-associated gene network that contains 46 genes that are mapped to Class IX of the WGCNA-identified co-expression clusters, predominantly in Cluster 19 (Fig. 5). Moreover, 17 of these 46 cuticular wax-associated genes are also hub genes for co-expression Clusters 1, 9, 18 and 19, and most of these hub genes are annotated as being involved in either “Lipid metabolism” or “Cellular respiration”.

Finally, the 150 genes that defined the network that encompasses both the cutin/suberin monomers and cuticular waxes, parsed into two subnetworks that predominantly reside within the WGCNA-identified co-expression Clusters 1 and 19 (Fig. 4 and 5). Whereas the Cluster 1 sub-network is dominated by genes related to “Photosynthesis”, the Cluster 19 sub-network is composed of genes associated with “Protein homeostasis”, “Lipid metabolism”, “External stimuli response”, and “Vesicle trafficking” (Fig. 5). Notably, 7 out of the 97 hub genes identified by the WGCNA-approach that reside in this network are components of the ubiquitin-26S proteasome system that mediates post-translational regulation of gene expression.

Validation of the conclusions reached from these complex analyses is gained by the fact that 16 genes residing in these gene networks have previously been experimentally confirmed to participate in cuticle deposition. These include genes that are either directly involved in cuticle deposition, or indirectly involved in this process by generating required precursors (Supplemental Table S8). For example, KCS1, KCS24 and KCR1 are components of the fatty acid elongation machinery that generates VLCFAs used for cuticular wax biosynthesis (Campbell et al., 2019); nsLTP2 and ECHINDA are involved in the transport of cuticular wax components to the extracellular space (Kim et al., 2012; McFarlane et al., 2014); a cytochrome P450 (CYP98A3) gene, GLOSSY14 and LACS1, each of which are required for cutin biosynthesis (Liu et al., 2021b); and the FDL1 transcription factor that acts as a positive regulator of cuticle biosynthesis (la Rocca et al., 2015; Castorina et al., 2020; Liu et al., 2021). Genes that are involved in the generation of cuticle precursors include acyl-carrier protein genes (MTACP2 and ACPT1) (Xia et al., 2009; Huang et al., 2017; Fu et al., 2020), three acyl-ACP desaturases (SACD1, SACD10 and SACD11) (Du et al., 2016; Zhao et al., 2019), and ATP-citrate lyase ACLA3 (Fatland et al., 2002). These genes are collectively involved in the generation of the precursors required for either *de novo* fatty acid biosynthesis or fatty acid elongation that ultimately generate the VLCFA precursors of the cuticular waxes.

The other members of these gene networks are primarily associated with two biological processes: a) photosynthesis and photosynthesis-related metabolism, and b) lipid metabolism (Fig. 6, Supplemental Fig. S7, Fig. S8 and Supplemental Table S8). Photosynthesis and photosynthesis-related metabolism genes include those encoding components of Photosystems I and II (e.g., LHCAs, LHCBs, PSAs and PSBs) (Teramoto et al., 2001; Bressan et al., 2018), enzymes for carbon fixation (e.g., GAPDHs and PEP1) (Yang et al., 2017), proteins maintaining and repairing photosynthetic machinery (e.g., chaperone proteins) (Bai et al., 2015), and proteins essential for chloroplast biogenesis (e.g., transcription factors G2 and GLK1) (Kobayashi and Masuda, 2016; Rocha et al., 2018; Fujii et al., 2019; Yeh et al., 2021) (Supplemental Table S8). Genes participating in lipid metabolism include those involved in triacylglycerol degradation (e.g., lipases SDP1 and MGAL5) (Kim et al., 2016; Fan et al., 2017), fatty acid β-oxidation (e.g., acyl-CoA dehydrogenase IBR3, acyl-CoA oxidase ACX4) (Kim and Miura, 2004), and phospholipid metabolism and signaling (e.g., diacylglycerol kinase DGK1) (Wang et al., 2006) (Supplemental Table S8). These latter findings indicate additional functionalities that likely indirectly impact cuticle deposition. Taken together, these findings demonstrate the complexity of the cuticle metabolome-associated gene networks, which consist of genes with confirmed roles in cuticle biosynthesis, potential cuticle-related genes, and those belonging to biological processes co-occurring with cuticle deposition.

**Figure 6.**
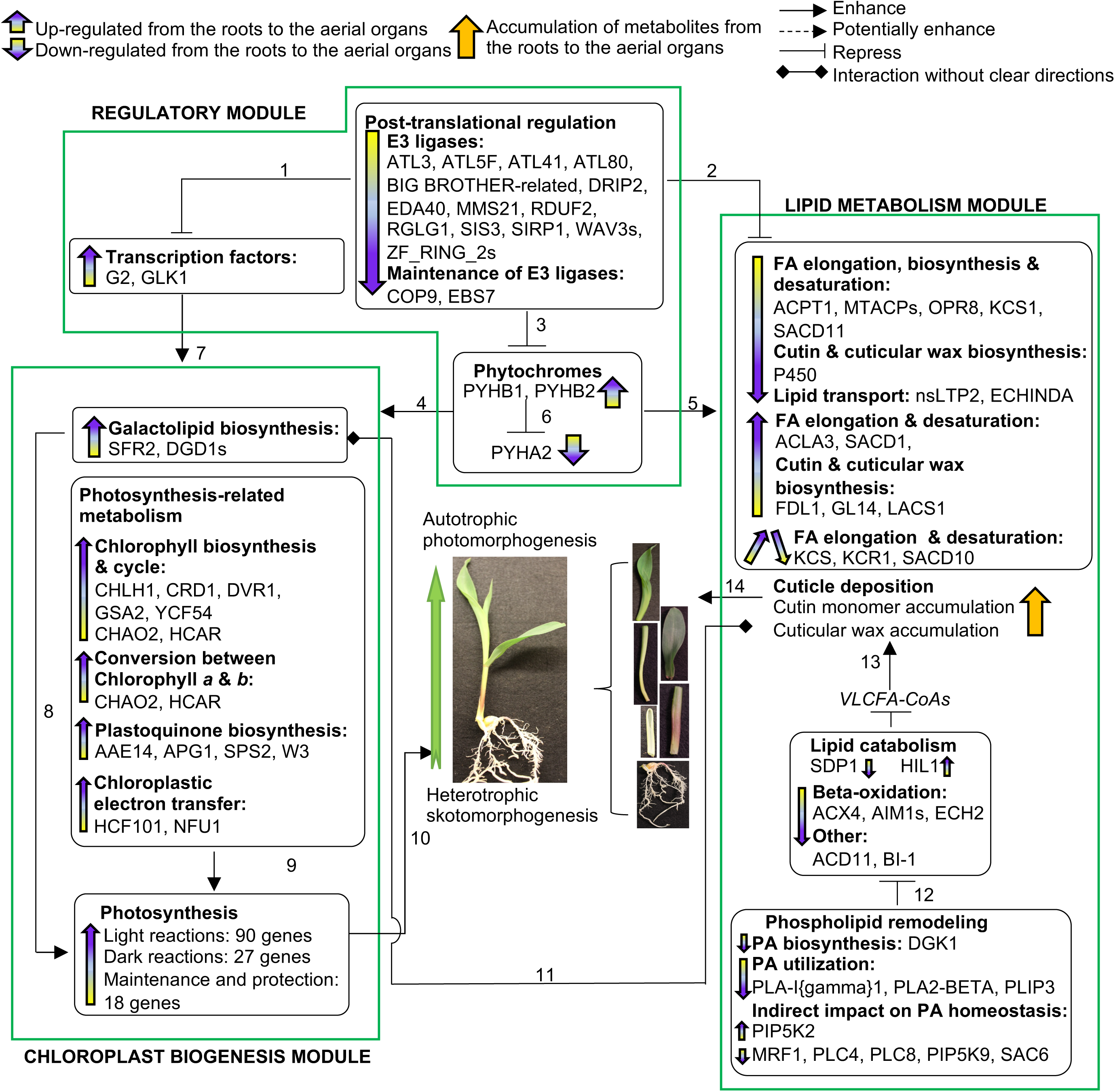
The regulatory program highlighting interactions between cuticle development and three functionality modules: a) regulatory module; b) lipid metabolism module and c) chloroplast biogenesis module. Numbered arrows identify interactions supported by literature based evidence: 1) Tokumaru et al., 2017; 2) Lü et al., 2012; Lu et al., 2013; Gil et al., 2017; Wang et al., 2018; Kim et al., 2019a; Liu et al., 2021a; Liu et al., 2021b; Wu et al., 2021; 3) Kim et al., 2017b; Han et al., 2020; 4) Legris et al., 2019; 5) Qiao et al., 2020; 6) Sheehan et al., 2007; 7) Kobayashi et al., 2014; Yeh et al., 2021; 8) Rocha et al., 2018; Fujii et al., 2019; 9) Schwenkert et al., 2010; Meguro et al., 2011; Wang et al., 2012; Reumann, 2013; Schlicke et al., 2014; Kim et al., 2015; Kobayashi and Masuda, 2016; Pattanayak and Tripathy, 2016; Kim et al., 2017a; Herbst et al., 2018; Dmitrieva et al., 2021; Satyanarayan et al., 2021; 10) Shi et al., 2020; 11) Xia et al, 2009; Wan et al., 2020; 12) Kim et al., 2019b; Cai et al., 2020; 13) Yeats and Rose, 2013; 14) Ingram and Nawrath, 2017. Supporting data including gene expression patterns can be found in Supplemental Fig. S7, Fig. S8, and Supplemental Table S8.

## DISCUSSION

The past few decades have manifested increasing challenges to maintain global food production. Although there are many causes to these challenges, a major contributor is global climate change that is the consequence of anthropogenic production of greenhouse gases (e.g., CO_2_, CH_4_, N_2_O and fluorinated gases), which are the aftermath of increased industrialization of human activities (Zaman et al., 2021). Projecting forward therefore, food crops will likely have to contend with increased abiotic and biotic stresses, including drought, salinity, solar radiation, elevated temperatures, pathogens and pests (Gong et al., 2014). As a water barrier the hydrophobic plant cuticle on aerial tissues (Riederer and Schreiber, 2001) and the suberin layer in the root periderm and endoderm (Hose et al., 2001; Suresh et al., 2022) will be integral in protecting plants from these increasing stresses, particularly in maintaining water status. These stresses may be analogous to the stresses that would have provided the selection pressure that led to the evolutionary appearance of the cuticle during the terrestrial colonization by land plants (Seale, 2020). In this context therefore, it is of high significance to have detailed, integrated physiological, genetic and biochemical insights of the processes that regulate the deposition of different cuticles by crop plants.

Global agriculture is highly dependent on cereal crops, with maize being ranked third, after wheat and rice (Cooper et al., 2014). As a biological system to study the cuticle, maize seedlings offer many advantages, including the availability of *glossy* mutants, first identified in 1928 (Hayes and Brewbaker, 1928), that affect the normal deposition of this material. Subsequent genetic and biochemical characterizations identified nearly 30 such *glossy* alleles (Schnable et al., 1994), and many of these loci have been molecularly characterized (Moose and Sisco, 1994; Hansen et al., 1997; Dietrich et al., 2005; Lauter et al., 2005; Sturaro et al., 2005; Liu et al., 2009; Li et al., 2013; Zheng et al., 2018; Li et al., 2019b; Alexander et al., 2020). The parallel application of similar strategies with mutant collections in other species, such as Arabidopsis (Kunst and Samuels, 2009) and tomato (Petit et al., 2021), have expanded the knowledgebase of the mechanisms of cuticle deposition. These studies have by and large validated the bifurcated structure of the metabolic process of cuticular wax deposition, first suggested by the characterization of maize mutants (Bianchi et al., 1985; Post-Beittenmiller, 1996). Namely, VLCFA products, generated by an ER localized fatty acid elongase, are further metabolized either through a reductive pathway or a decarbonylative pathway, collectively having the capacity to generate ∼500 different cuticular wax components.

More recently, whole genome research approaches have been employed to characterize the genetic architecture underlying cuticle deposition and function, broadening our knowledge of the genetic networks beyond the *glossy* genes. For example, gene co-expression networks in *glossy* mutants (Zheng et al., 2018) have revealed aspects of transcriptional regulation of cuticle biosynthesis, and other studies have queried the regulation of cuticle deposition by specific transcription factors (Castorina et al., 2020; Qiao et al., 2020; Yang et al., 2022). Genome- and transcriptome-wide association studies (Lin et al., 2020; Lin et al., 2022) have further probed the genetic underpinnings and regulation of cuticular conductance. Even with these findings, the complete genetic network underlying cuticular wax biosynthesis and its regulation remains undefined. Knowledge concerning the genetic network underlying cutin and suberin component of the cuticle is even less, primarily because of the lack of mutants that affect the assembly of these lipid polymers.

In this study, the cuticle metabolome was analyzed as the combination of the solvent extractable cuticular waxes, and the depolymerized products of the cutin (from aerial organs) or suberin (from roots) polymers. On a mole basis, at this early stage of seedling growth the cuticle of the aerial organs is composed primarily of cutin, whereas cuticular waxes account for between 10% and 25% of the cuticle. However, when one considers only the acyl constituents of cutin, the difference in relative abundance is less, with cuticular waxes accounting for 30-50% of the cuticle. In roots the cutin analog is suberin, and this polymeric material occurs at nearly 10-fold higher levels than the cuticular wax metabolites, which is consistent with the finding that on this organ the extracellular cuticle is constrained to the root cap (Berhin et al., 2019). These analyses revealed cuticle compositional differences, and these are primarily determined by organ differentiation. Indeed, high resolution spectroscopic imaging of the cuticle has revealed architectural changes during cuticle development, associated with both the cuticular waxes and the cutin components (Jun et al., 2010; Reynoud et al., 2021; Reynoud et al., 2022). One of the limitations associated with understanding the functional consequences of these alterations of the plant cuticle is the lack of detailed structural information concerning this hydrophobic matrix. This is particularly the case for the polymeric organization and structure of cutin and related polymers (i.e., suberin and sporopollenin), although new methods have recently been applied that are starting to reveal the molecular architecture of these biomaterials (Li et al., 2019a).

We used a systems-based strategy to ultimately explore the relationship between the cuticle metabolomes and the transcriptomes of maize seedling organs and identified gene networks associated with different cuticle metabolomes. Discerning biological relevance from complex multi-omics datasets can be challenging because large numbers of molecular features are being altered by the changes in the biological status of the system. This is primarily because phenotype datasets are relatively small as compared to the potential underlying genetic networks that determine the phenotype. For example, in this study, we investigated the relationships between cuticle composition (the phenotypes), measured as differences in the levels of ∼100 metabolites, and differences in the genetic expression program, measured as the accumulation levels of ∼30,000 transcripts, in the context of 24 different biological states (i.e., six different organs of four different maize genotypes). This complexity was harnessed by the application of two parallel statistical analytical pipelines, WGCNA-RF and a multi-omics integration pipeline, which reduced the coincidental associations between the metabolome and transcriptome. Genes that were identified by the former pipeline were classified as hub genes (821 genes), and genes identified by the latter pipeline were identified as putative cuticle-related genes (∼315 genes).

Confirmation of the validity of these two statistical pipelines to identify genes that are involved in establishing the cuticle is provided by two findings. First, both pipelines identify genes that have previously been experimentally confirmed to be directly involved in cuticle biosynthesis (Fig. 6). These include genes encoding functionalities needed for cuticular wax biosynthesis and deposition, including transcriptional regulation of the process, generation of upstream precursors to cuticle biosynthetic pathways, elongation of VLCFAs, and enzymatic components of the decarbonylative and reductive pathways of cuticular wax synthesis (Fatland et al., 2002; Fatland et al., 2005; Li-Beisson et al., 2010; McFarlane et al., 2014; la Rocca et al., 2015; Campbell et al., 2019; Castorina et al., 2020; Liu et al., 2021). In addition, four putative cuticle genes have also been identified as potentially involved in cuticle biosynthesis by genome- and transcriptome-wide association studies due to their role in cell wall biosynthesis (PAE5), vesicle trafficking (Ypt/Rab-GAPs), and regulation of cuticle development (WRKY71) (Lin et al., 2022). Moreover, in a gene co-expression network study based on >300 maize transcriptome datatsets, seven *glossy* genes known to regulate cuticle deposition were classified into the same co-expression cluster (Zheng et al., 2018). In the current study, these *glossy* genes also exhibited similarity in expression being more expressed in the encased leaves than the other seeding organs, though they did sort into different co-expression clusters.

Secondly, three gene networks that correspond to either cuticular wax biosynthesis, cutin/suberin biosynthesis, or a combination of the two, were deduced by integrating the outcomes from the two statistical pipelines. These three gene networks identify modules that are schematically illustrated in Figure 6. These modules are associated with a) chloroplast biogenesis, which include processes that incorporate galactolipid biosynthesis, chlorophyll metabolism and photosynthesis; b) regulatory functions, which include processes that incorporate transcription factors, post-translational regulation and phytohormones; and c) lipid metabolism, which include processes that incorporate cuticle generation, lipid catabolism and phospholipid remodeling.

These findings suggest the involvement of these processes in the generation of the seedling cuticle. However, because early seedling establishment involves the developmental transition from skotomorphogenesis to photomorphogenesis, many metabolic changes are associated with the transition from heterotrophic to autotrophic growth. For example, during this developmental transition there is a large increase in photosynthetic capability by the aerial organs of the seedling, as evidenced by the increased expression of genes belonging to chloroplast biogenesis processes (Fig. 6) that are essential for establishment and maintenance of photosynthetic machinery in the aerial organs. Concomitantly, there is down-regulation of genes involved in the lipid metabolism (Fig. 6) that mobilizes seed storage neutral lipids (i.e., triacylglycerol) and of fatty acid β-oxidation, which supported the initial skotomorphogenic growth prior to the acquisition of photosynthetic capability by the emerging seedling. Importantly, many of these processes that are associated with the skotomorphogenesis-photomorphogenesis transition have also been shown to be directly involved in the processes that establish the seedling cuticle. For example, cuticular wax mutants in different plant species exhibit changes in the biosynthesis of galactolipids, which are chloroplast-specific membrane lipids and thereby affect chloroplast development. Specifically, *glossy* mutants of navel orange affect the reallocation of carbon flux from cuticular waxes to chloroplast galactolipids (Wan et al., 2020). In Arabidopsis, mutation in the acyl-carrier-protein gene, *ACP4*, an ortholog of a cutin/suberin monomer-associated gene, *ACPT1*, causes abnormal cuticle formation and severe reduction in galactolipid accumulation (Xia et al., 2009). These findings substantiate our findings of interactions between cuticle biosynthesis and galactolipid biosynthesis during plastid development.

These analyses indicate another connection between photosynthesis and cuticle formation, namely phytochrome signaling, as a component of the regulatory module (Fig. 6). However, because phytochrome signaling stimulates the transition of seedling development from skotomorphogenesis to photomorphogenesis (Legris et al., 2019), it’s unclear how direct is the connection between photosynthesis and cuticle formation. Evidence that is supportive of this connection is the finding of altered cuticular wax compositions in maize phytochrome mutants, specifically *phyB1 phyB2* and *phyA1 phyA2* double mutants, and in the *phy* mutants of the moss *Physcomitrella patens*, suggesting that phytochrome-mediated light signaling regulates cuticle development (Qiao et al., 2020).

Apart from the phytochromes, the regulatory module associated with the transition from skotomorphogenesis to photomorphogenesis also includes twenty E3 ligase-coding genes that encode the key component of the ubiquitin-26S proteasome system, which is responsible for targeted protein degradation (Vierstra, 2009). Extensive prior studies (Kim et al., 2017; Han et al., 2020) have demonstrated that these E3 ligases are key regulators that maintain plant growth during extended periods of darkness, and this is achieved by the repression of photomorphogenesis mediated by the abundance of photoreceptors (e.g., phytochromes) and photomorphogenic transcription factors. Indeed, targeted protein degradation has already been associated with cuticle metabolism. For example, E3 ligases disrupt cuticle deposition by targeting degradation of cuticular wax biosynthetic proteins or transcription factors that activate cuticular wax biosynthesis (Lü et al., 2012; Gil et al., 2017; Kim et al., 2019a; Liu et al., 2021a; Wu et al., 2021; Wang et al., 2018; Lu et al., 2013). Additionally, the E3 ligase, SIS3, appears to be negatively regulated by the cuticle-controlling transcription factor FDL1 as demonstrated in a whole-genome scale identification of genes controlled by FDL1 (Liu et al., 2021). This finding is consistent with our observation that *Fdl1* shows an opposite expression level to that of *Sis3*. In addition to E3 ligases, we identified other cutin- and cuticular wax-related genes within the regulatory module that are associated with protein degradation, namely two proteins that interact with E3 ligases (i.e., COP9 and EBS7), and these control leaf epidermis development and cuticle formation (Liu et al., 2015b; Xie et al., 2020; Wu et al., 2021).

## CONCLUSION

Using a suite of approaches for joint metabolome-transcriptome analysis and gene co-expression network analysis, this study has identified gene networks associated with cuticle metabolome compositional changes in six maize seedling organs during the transition from skotomorphogenesis to photomorphogenesis. These gene networks consist of genes with experimentally confirmed roles in cuticle deposition and putative cuticle-related genes, and importantly identify new suites of genes that either directly or indirectly impact cuticle formation during seedling development. The gene networks identified herein provide new gene targets for improving the defense properties of the cuticle, thereby better arming fledgling seedlings to withstand the new challenges to crop productivity.

## METHODS

### Plant materials

Seeds of the maize inbred lines, B73, Mo17, and reciprocal hybrids, B73×Mo17 and Mo17×B73 were sown in flats of 16 pots filled with Redi-Earth soil (Hummert, USA). Three biological replicates were grown per genotype, with each replicate grown in its own flat that included 8 individual pots (1 seed per pot) for each of the four genotypes. Plant were watered every 48 h, with a solution supplemented with 220 ppm calcium nitrate (Fisher, USA) and 96 ppm magnesium sulfate (Fisher, USA). In addition, the soil was treated with Marathon pesticide (Hummert, USA) to control pests. Plants were grown in a climate-controlled greenhouse, maintained at 30% humidity, under a diurnal cycle of 16 h of illumination (at light intensity 230 µE m^-2^ s^-1^) and 8 h of darkness, at 28 °C and 21 °C, respectively. Plants were harvested for organ dissection based on their height (between 12 cm and 15 cm). Generally, organs from 5-6 plants per replicate per genotype were sampled and pooled for RNA and cuticle extraction. Coleoptiles from B73 and Mo17 were harvested at 6 days after sowing, and at 5 days after sowing for the reciprocal hybrids. The other seedling organs (roots, sheath of first leaf, 1^st^ leaf blade, encased leaves, and 2^nd^ leaf blade) were harvested from inbreds B73 and Mo17 at nine days after sowing, and at eight days after sowing from the reciprocal hybrids. Pools of each of these tissues were flash frozen in liquid nitrogen and stored at -80 °C.

### Cuticular wax extraction and quantification

Pooled tissue samples were separately dipped for 30 s at room temperature into 10 ml of chloroform containing 5 μg of heptadecanoic acid (Sigma, St. Louis) as the internal quantification standard. The extracts were dried under a stream of nitrogen gas and derivatized by incubation at 70 °C for 30 min with 100 μl of N,O–Bis (trimethylsilyl) trifluoroacetamide (BSTFA) and 100 μl anhydrous pyridine (Sigma, St. Louis). After the derivatization reaction, excess BSTFA and pyridine were evaporated under a stream of nitrogen gas, and the residue was dissolved in 500 μl of heptane:toluene (1:1, v/v) and subjected to gas chromatography-mass spectrometric (GC-MS) analysis. Cuticular wax metabolites were identified using AMDIS software (Stein, 1999) with the NIST14 mass spectral library (http://nistmassspeclibrary.com) and metabolite concentrations were calculated relative to the internal standard and presented as µmol.g^-1^ dry weight.

### Cutin and suberin monomer extraction and quantification

Cutin from the aerial organs, and suberin from root samples were extracted in parallel from aliquots of the same pooled tissue samples as those used for cuticular wax analysis. Dried 10-20 mg of tissue was exhaustively delipidated by immersing overnight at room temperature in isopropanol containing 0.01% (w/v) butylated hydroxytoluene (BHT) in PTFE screw-capped glass tubes on a rotatory shaker. The solvent was discarded the next day, and the tissue was extracted with two aliquots of chloroform: methanol (2:1) containing 0.01% (w/v) BHT, and subsequently with an aliquot of methanol containing 0.01% (w/v) BHT. Each of these extractions was for a period of 2-h at room temperature. Finally, the methanol was discarded and the delipidated residue preparation was dried by lyophilization to a constant dry weight.

Acid-catalyzed transmethylation was conducted by adding to the dry residue, 2 ml of freshly prepared methanolic sulfuric acid (4% v/v) and 0.2 ml of toluene, along with 10 μg of heptadecanoic acid as an internal quantification standard. The mixture was heated at 80 °C for 2 h, and after cooling, the mixture was extracted with 4 mL dichloromethane and 1 mL of 0.9% (w/v) NaCl in 100 mM Tris-HCl buffer, pH 8.0. The non-polar phase was separated from the polar phase by centrifugation at 1500 rpm for 2 min, and the two phases were recovered separately and dried. Each sample was silylated at 70 °C for 30 min, with 100 μl anhydrous pyridine and 100 μl BSTFA. The mixture was subsequently dried under a stream of nitrogen gas, and the residue was dissolved in 500 μl of heptane:toluene (1:1, v/v) and subjected to GC-MS analysis (Bhunia et al., 2018).

Methylated and silylated cuticular wax constituents and cutin/suberin monomers were analyzed using an Agilent Technologies Model 7890A gas chromatograph coupled to Model 5975C mass spectrometer. GC conditions were as follows: the inlet temperature was held constant at 280 °C, the helium carrier gas was at a constant flow rate of 1 mL.min^−1^ through an Agilent 122-0112 DB-1ms column (15 m × 250 µm × 0.25 µm). The oven temperature was initially set on 70 °C, and then raised by 10 °C.min^−1^ to 340 °C and held at that temperature for 6 min; the transfer line was set to 280 °C.

Detection and quantification of individual monomers was accomplished using an Agilent Model 5975C mass spectrometer under standard conditions with 280 °C ion source. Absolute quantification was determined by comparing the ion signal of each peak to that of the heptadecanoic acid internal standard.

### RNA extraction and sequencing

RNA was extracted from pooled tissues by the TRIZOL method (Invitrogen, Carlsbad) per the manufacturer’s protocol and subsequently purified using a Qiagen RNeasy MinElute Cleanup Kit. RNA-seq libraries for each biosample were prepared using a KAPA Stranded RNA-Seq Library Preparation Kit (Illumina, San Diego, CA) and subjected to RNA-seq at the Beijing Genomics Institute (BGI) Americas. The 100-cycle paired-end sequencing data were acquired from nine lanes of Illumina HiSeq 4000, with 24 transcriptomes sequenced per lane. Each biological replicate of the six seedling organs from four genotypes was sequenced in three separate lanes to ensure appropriate sequencing depth.

Preprocessing of resultant RNA-sequencing reads for removal of adapter sequences and subsequent data quality control was performed by the FastQC function as described in McNinch et al. (2020), revealing an average read length of 100 bp and a median read depth of 39.1 million reads per biological replicate. Reads from all samples were aligned to the B73 genome (Jiao et al., 2017) that already incorporated Mo17 variants (Sun et al., 2018) into the genome build, prior to aligning by the HISAT2 alignment program (Kim et al., 2015). Subsequently, transcripts were assembled and quantified on a fragments per kb of transcript per million mapped reads (FPKM) basis as described in McNinch et al. (2020).

### Statistical analysis of cuticle metabolomes

The effects of seedling genotype, organ, and the genotype × organ interaction on the abundance of each lipid class of cutin/suberin monomers or cuticular waxes was evaluated by Type I Analysis of Variance (ANOVA). Post-hoc Tukey’s honest significant difference (HSD) test was applied to further examine the genotype differences within each organ. The ANOVA and HSD tests were conducted using JMP® (Version 15.0. SAS Institute Inc., Cary, NC, 1989–2021).

### PCA and tSNE visualization of metabolomics and transcriptomics data

Prior to PCA and tSNE visualization, metabolomics and transcriptomics datasets were transformed such that the average expression of individual genes or metabolites was zero, with the corresponding variance set to one. For each dataset, PCA was performed using princomp function in the R/stats package (R Core Team, 2019), and tSNE visualization, based on all principal components was performed using the Rtsne function in the R/Rtsne package (Krijthe, 2015).

### Gene co-expression network analysis

The co-expressed gene clusters were determined using R/WGCNA package (Langfelder and Horvath, 2008) as described in McNinch et al. (2020). The transcriptome datasets with the expression profiles of 30,931 genes were first subjected to data filtering with the goodSampleGenes function, which left 22,841 genes that are expressed in more than half of the samples. Briefly, an adjacency matrix among the expressed genes was first derived from biweight mid-correlations with a soft-threshold power of 16. A topographical overlap matrix was next constructed based on the adjacency matrix and used for the hierarchical clustering conducted using hclust. Co-expressed gene clusters were refined from the resultant dendrogram using function cutreeDynamic followed by mergeCloseModules. An eigengene value was computed for each cluster to represent the overall expression pattern of all genes in the cluster. A second hierarchical clustering and Pearson correlation analysis was performed among the cluster eigengenes to classify clusters with similar expression patterns into separate classes. Each class was comprised of clusters with correlation coefficients greater than 0.5.

Finally, hub genes were identified for each co-expression cluster. A gene is considered as a cluster hub gene if it satisfies the following three requirements: (1) an above average significant connection degree (Das et al., 2017), i.e., it is connected in its expression pattern with a large number of other genes in the cluster; (2) the correlation value between the candidate hub gene and the corresponding eigengene is >0.8; and (3) the correlation between the candidate hub gene and traits of interest, in this case tSNE components for metabolomics data, is greater than an assigned cutoff value (Liu et al., 2019). Different correlation cutoffs were selected for each cluster such that ∼5% of the genes in a cluster were identified as hub genes.

### Multi-omics integration pipeline: gene-to-metabolite associations

Partial Least Square regression (PLS), sparse Partial Least Square regression (sPLS) and random generalized linear model (rGLM) were used to identify cuticle-related genes having different levels of expression in the different seedling organ samples. Data centering was applied in PLS to make the average of individual metabolite concentration or gene expression to be zero. Data scaling for sPLS and rGLM was performed so that the variance of metabolite concentration or gene expression was one. PLS and sPLS were conducted using scripts that were implemented in R language, and rGLM conducted using the R/randomGLM package (Song et al., 2013), with modifications to the original scripts (https://github.com/ketingchen/MultiOmicsIntegration.git).

Two types of response variables were independently used to represent the compositional changes in the cuticle metabolome: (1) the concentration profiles of all metabolites (PLS and sPLS), and (2) single-value tSNE components that captured the major variations across metabolomes. PLS and sPLS statistical methods utilized the former type of responses, while rGLM used the latter, because this method can only incorporate a single-value response variable.

In each of these models, an importance score, *S*, was calculated to represent the contribution of individual genes in predicting metabolome composition. The probability of obtaining a background importance score, *S*_B_, exceeding or equaling *S* (*P*(*S*_B_ ≥ *S*)) in datasets devoid of transcriptome-metabolome correlations was evaluated by a permutation test that is described in the Supplemental Methods . A gene is selected as a putative cuticle-related gene by a model when *S*>1 and *P*(*S_B_* ≥ *S*)<0.01.

This suite of multivariate statistical methods was applied separately to interrogate the gene-to-cutin/suberin monomer and gene-to-cuticular wax constituent associations. An individual expressed gene was considered a putative cuticle-related gene impacting cutin monomer and/or cuticular wax composition if it was selected by at least one of the three statistical approaches.

### Multi-omics integration pipeline: gene cluster-to-metabolite associations

The association between a co-expressed gene cluster and cuticle metabolomes was interrogated by a random forest regression model that incorporated the eigengene expression for each cluster and the expression data of non-clustered individual genes as the predictors, and the tSNE components for cutin/suberin monomers or cuticular waxes as the response variables. Three metrics were used to evaluate the importance of a gene cluster (or an individual gene) on the metabolome compositions: (1) predictive R^2^ (i.e., 1 – sum of squares error/total sum of squares), (2) root mean square error, and (3) mean absolute percent error (100 × | predicted value – actual value | /actual value). In detail, for each predictor, two random forest models were constructed, with one including the full set of predictors, and the other having one specific predictor removed. The difference in each evaluating metric was subsequently calculated between the two models. A bootstrapping strategy was used to calculate the p-values associated with the difference, which were later corrected among all predictors to control the false discovery rate <5%. A predictor was significantly associated with the compositional changes of cutin monomers and/or cuticular waxes if removal of this predictor from the random forest model reduced the model performance in at least two metrics, with p-values <0.01. The Random Forest model was performed by R/randomForest package (Liaw and Wiener, 2002) and the bootstrapping strategy that evaluates the predictor importance and computes the p-values, was performed using scripts that were implemented in R language (https://github.com/ketingchen/MultiOmicsIntegration.git).

### Gene functional enrichment analysis

GO enrichment tests for the biological process domain were performed using the R/topGO package (Alexa and Rahnenfuhrer, 2021) and maize-GAMER GO annotations (Wimalanathan et al., 2018), as described in McNinch et al. (2020). The resultant p-values were corrected to control the false discovery rate < 5%. GO terms with corrected p-values <0.0001 were considered as significantly enriched. The significant GO terms were then trimmed by REVIGO (Supek et al., 2011), a web server tool that removes functional redundancy of each list of GO terms via the identification of the most informative representative terms according to “parent-child” hierarchy and semantic similarity among GO terms.

Annotations of MapMan pathway categories (i.e., function bins) for all protein-coding maize genes were derived from the maize B73 draft genome version 4, as retrieved from the MapMan website (https://mapman.gabipd.org/mapman). MapMan-based function enrichment tests were performed by Fisher’s exact test. The resultant p-values were corrected to control the false discovery rate < 5%. MapMan bins with corrected p-values <0.05 were considered as significantly enriched. The putative cuticle-related genes identified by multi-omics integration analysis and annotated with significantly enriched GO terms and MapMan bins were visualized in Cytoscape (version 3.8.2) (Shannon et al., 2003) to present their co-expression clustering membership that was determined by WGCNA.

### Accession Numbers

RNAseq data and the associated biosample metadata are available in the NCBI Sequence Read Archive, accession PRJNA903105.

## ACKNOWLEDGEMENTS

We thank Dr. Liza E. Alexander for assistance with sample preparation and cuticle extraction, Dr. Xuefeng Zhao for initial assistance in RNA-Seq data analyses, Dr. Ann Perera and Dr. Lucas Showman of the W. M. Keck Metabolomics Research Laboratory for assistance and advice regarding metabolomic profiling, and Dr. Dan Nettleton and Dr. Nick Lauter for advice on transcriptome analysis. This research was supported by NSF grants IOS-1354799 and MCB-2212799 to BJN and MDY-N, and NSF grant NSF-PGRP-1546727 to MDY-N. Additional support was provided by NIFA-USDA Hatch project IOW03649 (MDY-N), NIFA-USDA Multi-state Project IOW05660 (BJN and MDY-N) and through Iowa State University’s Center for Metabolic Biology. Mention of trade names or commercial products in this publication is solely for the purpose of providing specific information and does not imply recommendation or endorsement by the USDA. USDA is an equal opportunity provider and employer. The content of this paper is however solely the responsibility of the authors and does not represent the official views of the NSF, NIFA or USDA.

## FIGURE LEGENDS

### SUPPLEMENTAL MATERIAL

**Supplemental Table S1**: Cutin/suberin monomer and cuticular wax constituents profiled from maize seedlings of B73, Mo17, and the reciprocal hybrids. Provided as an external file.

**Supplemental Table S2:** Co-expressed gene cluster and the corresponding hub genes determined by WGCNA, and gene cluster-to-metabolome association analysis by random forest regression model. Provided as an external file.

**Supplemental Table S3:** Putative cuticle-related genes associated with the compositional changes of cutin/suberin monomers and/or cuticular waxes as identified by multi-omics integration pipeline. Provided as an external file.

**Supplemental Table S4:** MapMan function enrichment test for putative cuticle-related genes that are only associated with cutin/suberin monomer composition (cutin/suber monomer-associated genes) or cuticular wax composition (cuticular wax-associated genes), or are associated with compositions of both cutin/suberin monomers and cuticular waxes (cutin- and cuticular wax-associated genes).

**Supplemental Table S5:** Gene ontology enrichment analysis for cutin/suberin monomer-associated genes. Provided as an external file.

**Supplemental Table S6:** Gene ontology enrichment analysis for cuticular wax-associated genes. Provided as an external file.

**Supplemental Table S7:** Gene ontology enrichment analysis for cutin/suberin- and cuticular-wax associated genes. Provided as an external file.

**Supplemental Table S8:** Putative cuticle-related genes that are annotated with significantly enriched metabolic processes that directly or indirectly interact with cuticle biosynthesis. Provided as an external file.

**Supplemental Figure S1:**
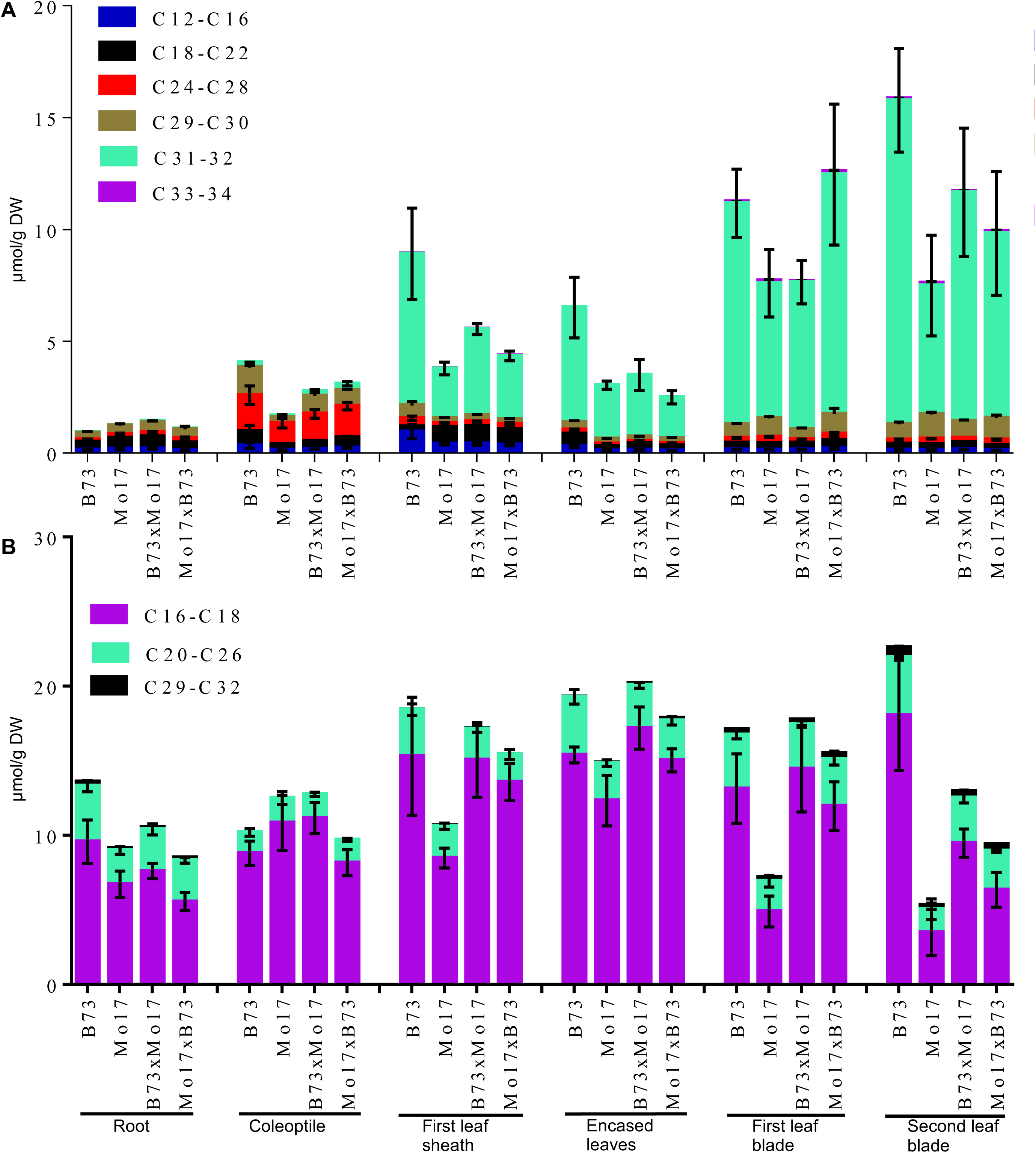
Comparison of the alkyl chain lengths of cuticular waxes (A), and cutin and suberin monomers (B) in maize seedlings.

**Supplemental Figure S2:**
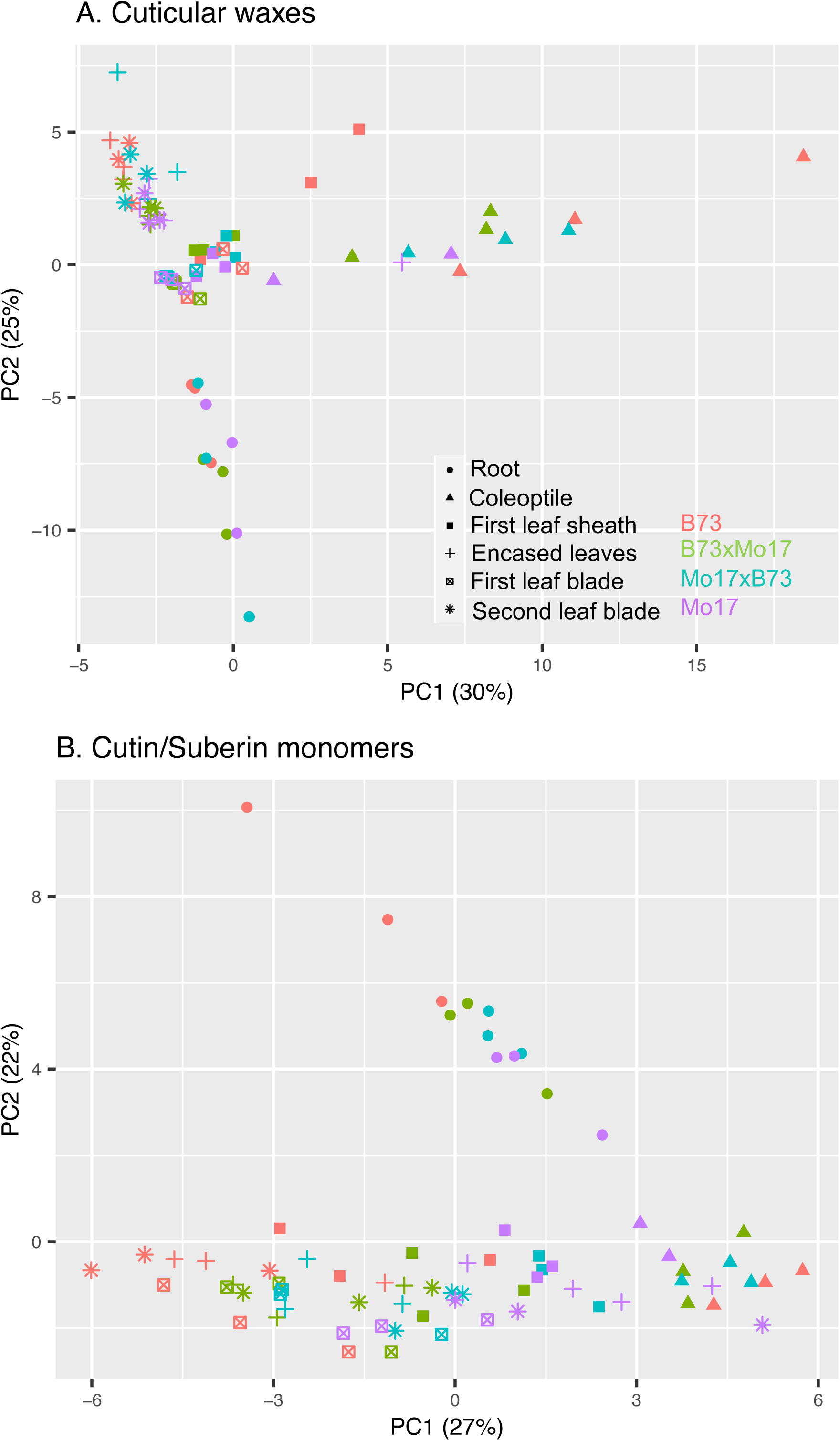
PCA for cuticular waxes (A) and cutin/suberin monomers (B). Symbol color denotes genotype and symbol shape denotes seedling organ. The percentages listed represents the percent of variance explained by that PC.

**Supplemental Figure S3:**
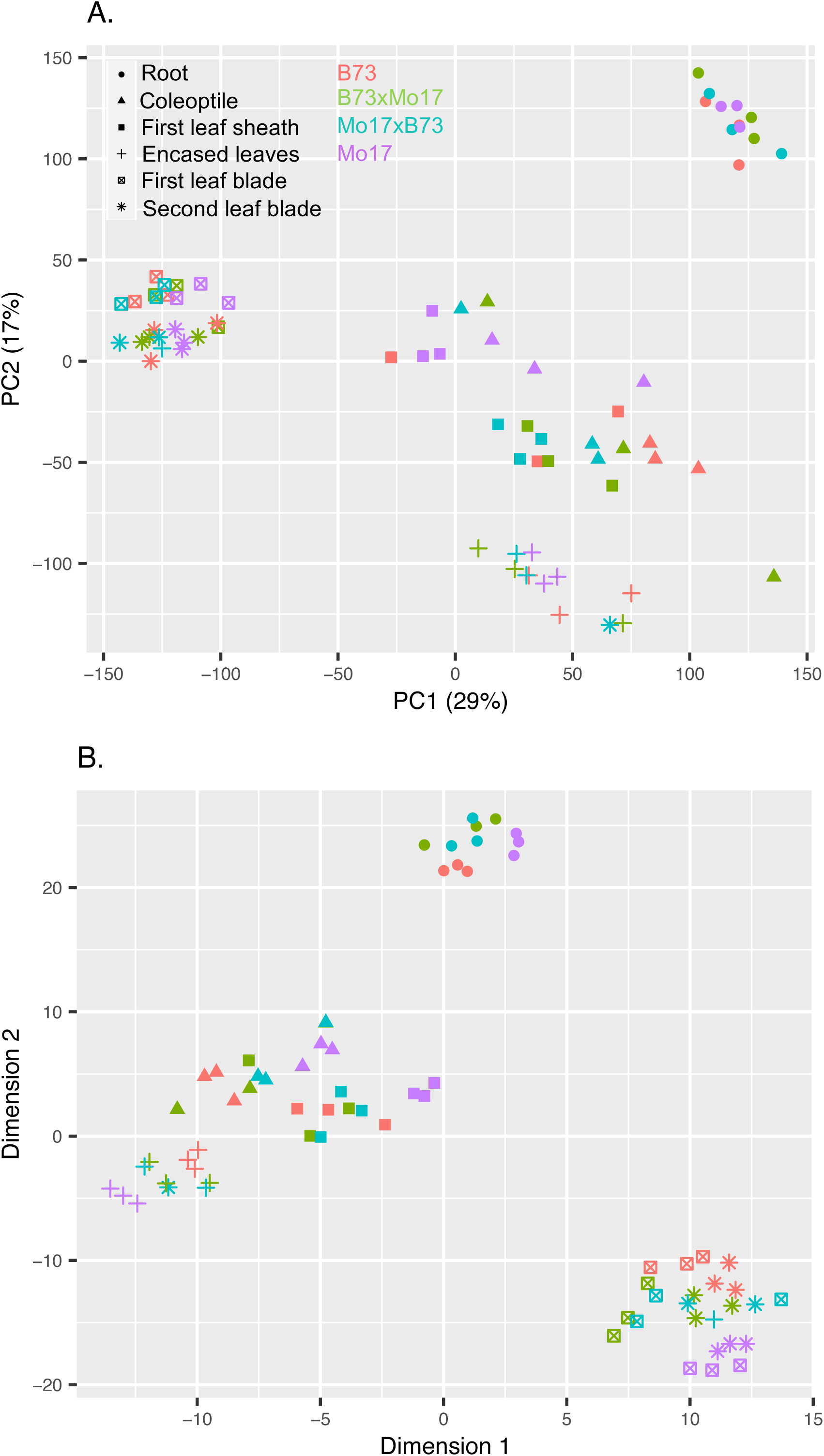
PCA (A) and tSNE (B) visualization for transcriptome datasets. The percentages listed in (A) represents the percent of variance explained by that PC.

**Supplemental Figure S4:**
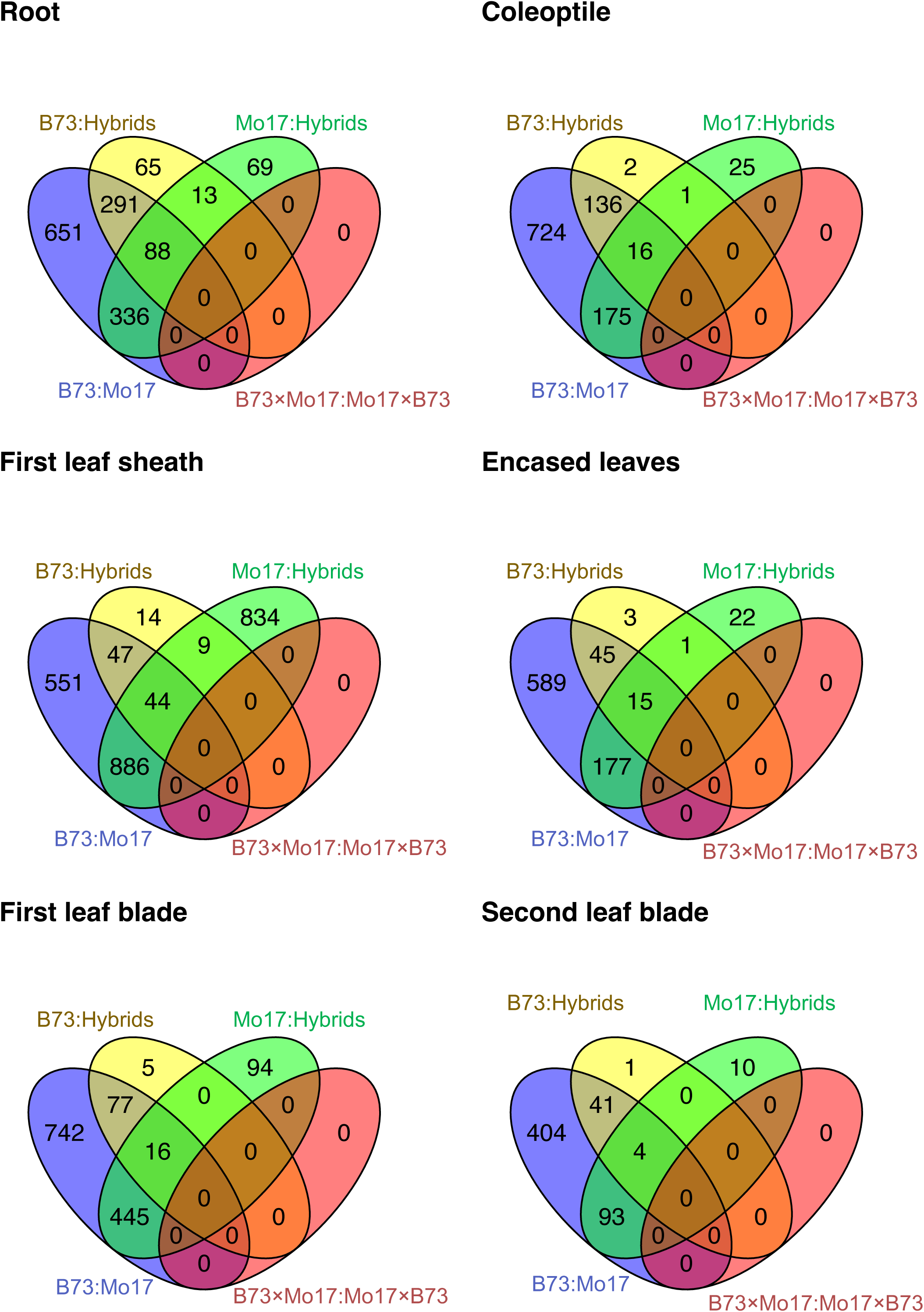
Venn diagram comparison of the differentially expressed genes between every pair of genotypes in each seedling organ.

**Supplemental Figure S5:**
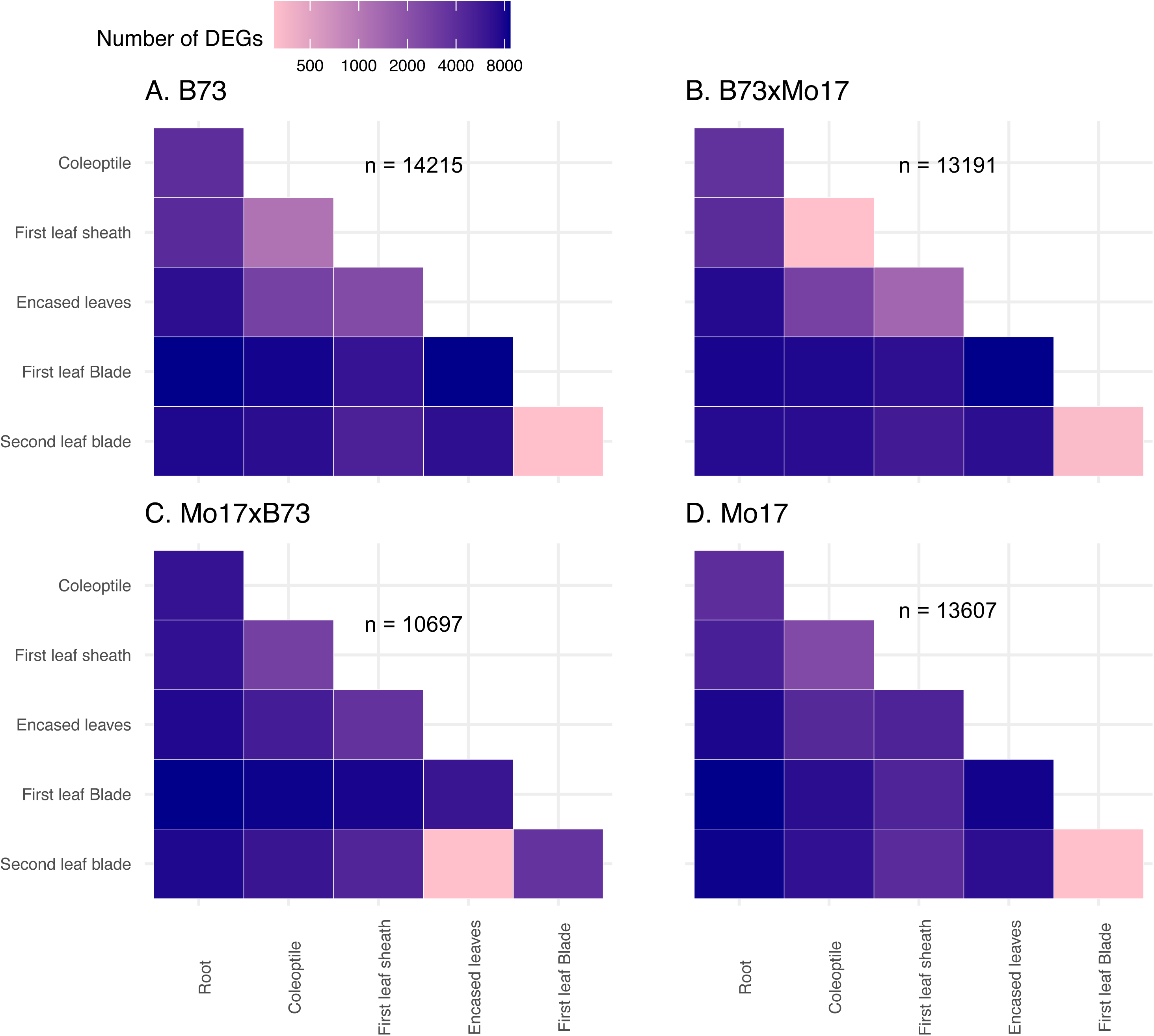
Heatmap representation of the number of differentially expressed genes between every pair of seedling organs in B73, Mo17, and the reciprocal hybrids. The total number of non-redundant differentially expression genes is indicated for each genotype (n).

**Supplemental Figure S6:**
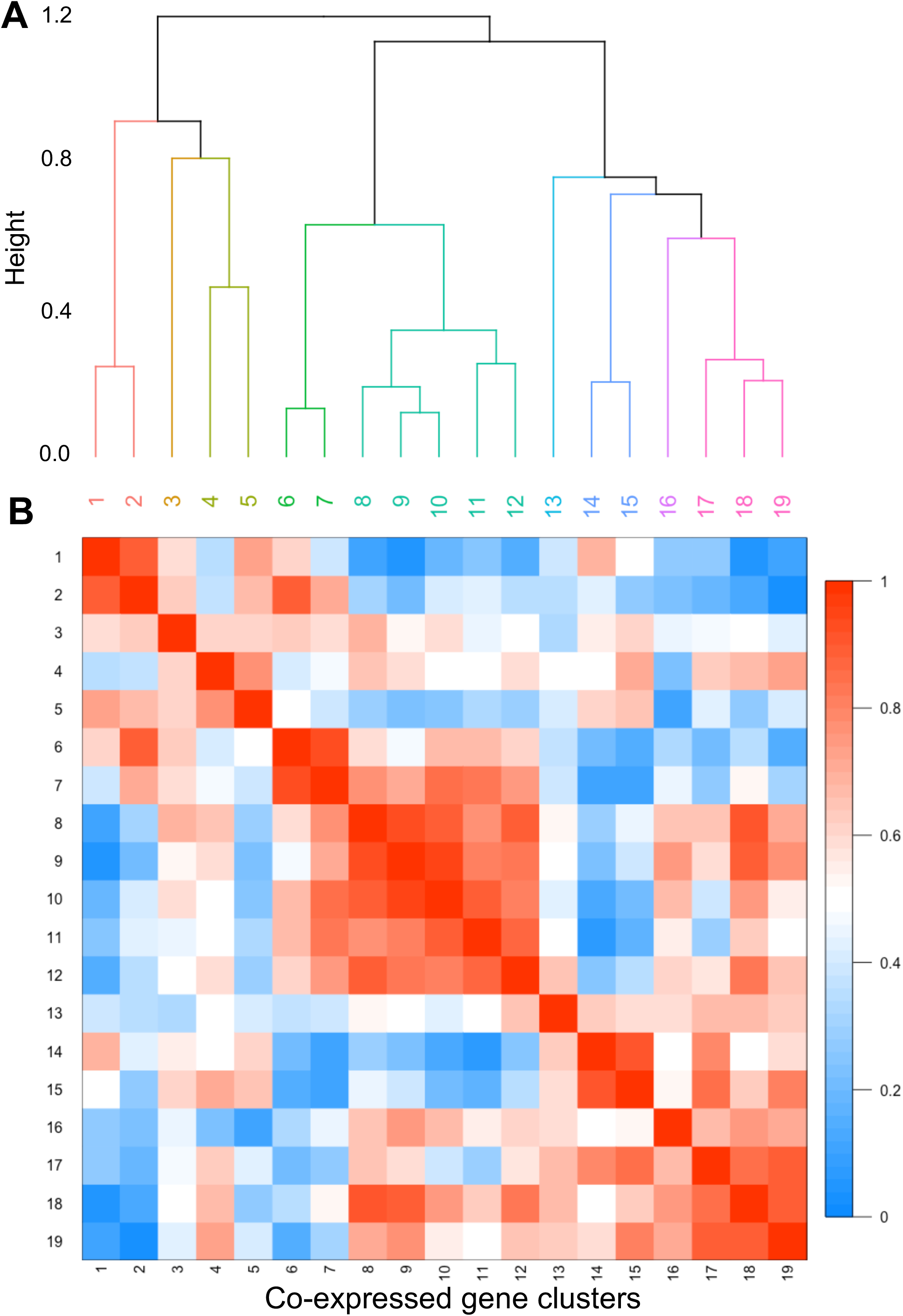
Similarity among eigengene expression for the 19 WGCNA-based co-expressed gene clusters (present in Fig. 4) as evaluated by hierarchical clustering (A) and Pearson-correlation based adjacency heatmap (B).

**Supplemental Figure S7:**
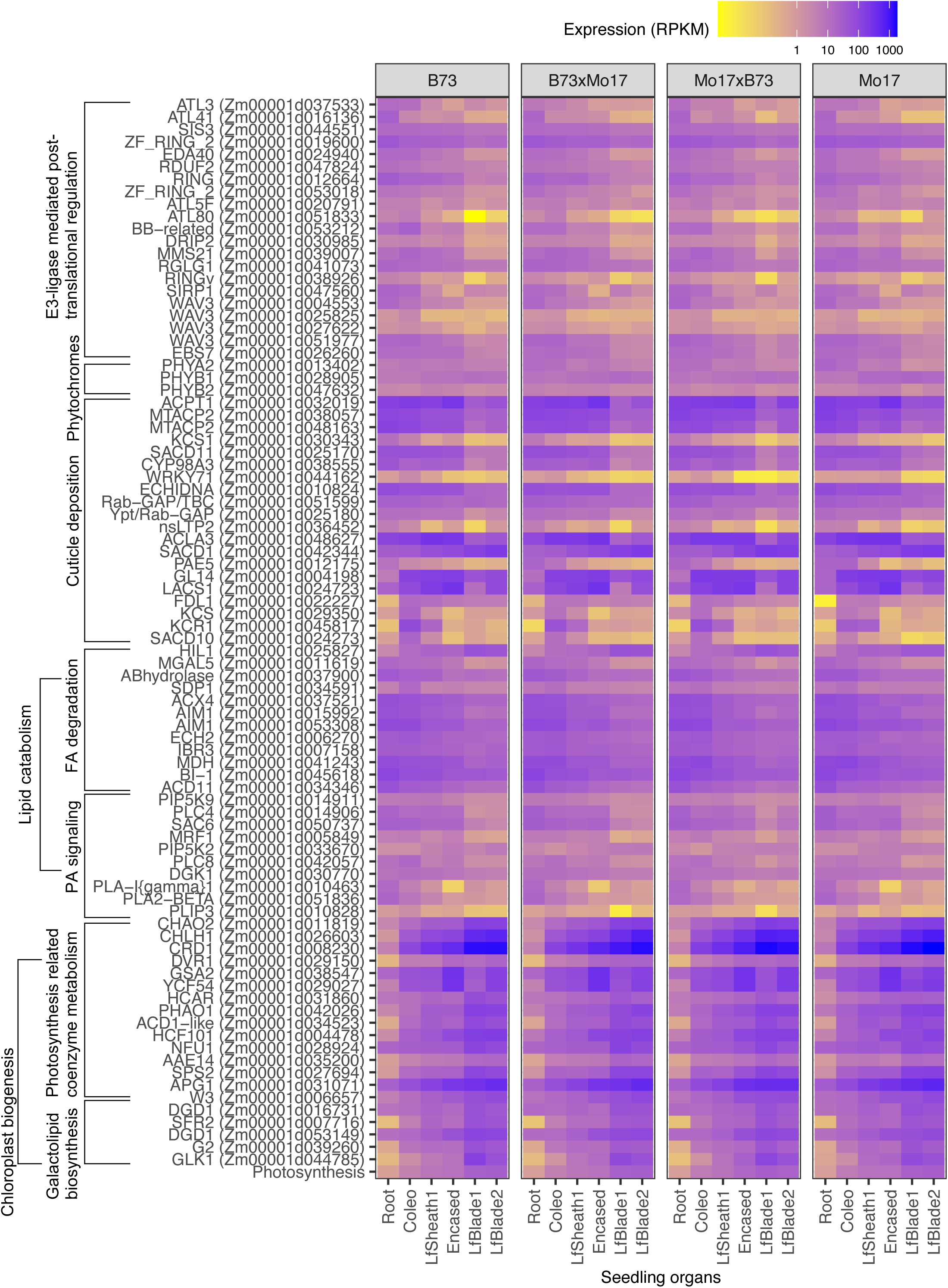
Heatmap visualization of the expression of the genes that were identified by the multi-omics integration analysis and were assigned to significantly enriched metabolic processes according to the enrichment analysis of GO terms and MapMan function bins. The expression of 135 photosynthesis genes is summarized and presented as a single eigengene, labeled “Photosynthesis” at the bottom of the plot. The expression of these photosynthesis genes are separately presented in Supplemental Fig. S8. Abbreviations: Coleo, coleoptile; LfSheath1, fist leaf sheath; Encased, encased leaves; LfBlade1, first leaf blade; LfBlade2, second leaf blade

**Supplemental Figure S8:**
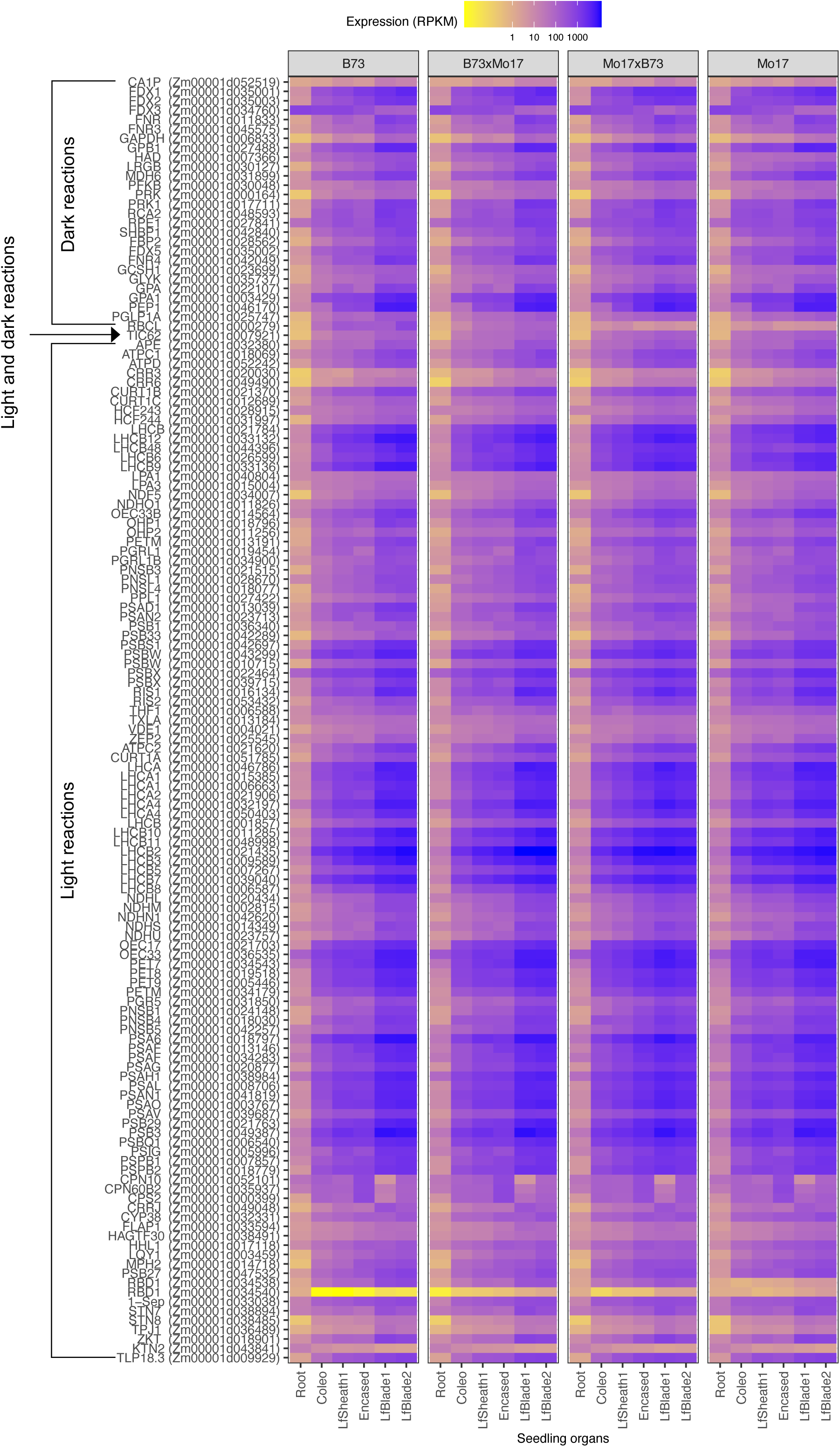
Heatmap visualization of the expression of 135 “Photosynthesis”-pathway associated genes as identified by MapMan analysis. Abbreviations: Coleo, coleoptile; LfSheath1, fist leaf sheath; Encased, encased leaves; LfBlade1, first leaf blade; LfBlade2, second leaf blade

## REFERENCES

Alexa A, Rahnenfuhrer J (2021) topGO: Enrichment analysis for gene ontology. R package version 2.46.0

Alexander LE, Okazaki Y, Schelling MA, Davis A, Zheng X, Rizhsky L, Yandeau-Nelson MD, Saito K, Nikolau BJ (2020) Maize *Glossy2* and *Glossy2-like* genes have overlapping and distinct functions in cuticular lipid deposition. Plant Physiol 183: 840–853

Ashburner M, Ball CA, Blake JA, Botstein D, Butler H, Cherry JM, Davis AP, Dolinski K, Dwight SS, Eppig JT, et al (2000) Gene Ontology: Tool for the unification of biology. Nat Genet 25: 25–29

Avato P, Bianchi G, Salamini F (1987) Ontogenetic variations in the chemical composition of maize surface lipids. The metabolism, structure, and function of plant lipids. Springer New York, Boston, MA, pp 549–551

Baxter I, Hosmani PS, Rus A, Lahner B, Borevitz JO, Muthukumar B, Mickelbart M v., Schreiber L, Franke RB, Salt DE (2009) Root suberin forms an extracellular barrier that affects water relations and mineral nutrition in Arabidopsis. PLoS Genet 5: e1000492

Berhin A, de Bellis D, Franke RB, Buono RA, Nowack MK, Nawrath C (2019) The root cap cuticle: A cell wall structure for seedling establishment and lateral root formation. Cell 176: 1367–1378.e8

Bhanot V, Fadanavis SV, Panwar J (2021) Revisiting the architecture, biosynthesis and functional aspects of the plant cuticle: There is more scope. Environ Exp Bot 183: 104364

Bianchi A, Bianchi G, Avato P, Salamini F (1985) Biosynthetic pathways of epicuticular wax of maize as assessed by mutation, light, plant-age and inhibitor studies. Maydica 30: 179–198

Bourgault R, Matschi S, Vasquez M, Qiao P, Sonntag A, Charlebois C, Mohammadi M, Scanlon MJ, Smith LG, Molina I (2020) Constructing functional cuticles: Analysis of relationships between cuticle lipid composition, ultrastructure and water barrier function in developing adult maize leaves. Ann Bot 125: 79–91

Bressan M, Bassi R, Dall’Osto L (2018) Light harvesting complex I is essential for Photosystem II photoprotection under variable light conditions in *Arabidopsis thaliana*. Environ Exp Bot 154: 89–98

Campbell AA, Stenback KE, Flyckt K, Hoang T, Perera MAD, Nikolau BJ (2019) A single-cell platform for reconstituting and characterizing fatty acid elongase component enzymes. PLoS One 14: e0213620

Carlson MR, Zhang B, Fang Z, Mischel PS, Horvath S, Nelson SF (2006) Gene connectivity, function, and sequence conservation: Predictions from modular yeast co-expression networks. BMC Genomics 7: 40

Castorina G, Domergue F, Chiara M, Zilio M, Persico M, Ricciardi V, Horner DS, Consonni G (2020) Drought-responsive *ZmFDL1/MYB94* regulates cuticle biosynthesis and cuticle-dependent leaf permeability. Plant Physiol 184: 266–282

Cooper M, Gho C, Leafgren R, Tang T, Messina C (2014) Breeding drought-tolerant maize hybrids for the US corn-belt: Discovery to product. J Exp Bot 65: 6191–6204

Das S, Meher PK, Rai A, Bhar LM, Mandal BN (2017) statistical approaches for gene selection, hub gene identification and module interaction in gene co-expression network analysis: An application to aluminum stress in soybean (*Glycine max* L.). PLoS One 12: e0169605

Dennison T, Qin W, Loneman DM, Condon SGF, Lauter N, Nikolau BJ, Yandeau-Nelson MD (2019) Genetic and environmental variation impact the cuticular hydrocarbon metabolome on the stigmatic surfaces of maize. BMC Plant Biol 19: 430

Dietrich CR, Perera MADN, D. Yandeau-Nelson M, Meeley RB, Nikolau BJ, Schnable PS (2005) Characterization of two GL8 paralogs reveals that the 3-ketoacyl reductase component of fatty acid elongase is essential for maize (*Zea mays* L.) development. Plant J 42: 844–861

Du H, Huang M, Hu J, Li J (2016) Modification of the fatty acid composition in Arabidopsis and maize seeds using a stearoyl-acyl carrier protein desaturase-1 (*ZmSAD1*) gene. BMC Plant Biol 16: 137

Fan J, Yu L, Xu C (2017) A central role for triacylglycerol in membrane lipid breakdown, fatty acid *β* -oxidation, and plant survival under extended darkness. Plant Physiol 174: 1517–1530

Fatland BL, Ke J, Anderson MD, Mentzen WI, Cui LW, Allred CC, Johnston JL, Nikolau BJ, Wurtele ES (2002) Molecular characterization of a heteromeric ATP-citrate lyase that generates cytosolic acetyl-Coenzyme A in Arabidopsis. Plant Physiol 130: 740–756

Fatland BL, Nikolau BJ, Wurtele ES (2005) Reverse genetic characterization of cytosolic acetyl-CoA generation by ATP-citrate lyase in Arabidopsis. Plant Cell 17: 182–203

Fich EA, Segerson NA, Rose JKC (2016) The plant polyester cutin: Biosynthesis, structure, and biological roles. Annu Rev Plant Biol 67: 207–233

Fu X, Guan X, Garlock R, Nikolau BJ (2020) Mitochondrial fatty acid synthase utilizes multiple acyl carrier protein isoforms. Plant Physiol 183: 547–557

Fujii S, Nagata N, Masuda T, Wada H, Kobayashi K (2019) Galactolipids are essential for internal membrane transformation during etioplast-to-chloroplast differentiation. Plant Cell Physiol 60: 1224–1238

Gong F, Yang L, Tai F, Hu X, Wang W (2014) “Omics” of maize stress response for sustainable food production: Opportunities and challenges. OMICS 18: 714–732

Han X, Huang X, Deng XW (2020) The photomorphogenic central repressor COP1: conservation and functional diversification during evolution. Plant Commun 1: 100044

Hansen JD, Pyee J, Xia Y, Wen TJ, Robertson DS, Kolattukudy PE, Nikolau BJ, Schnable PS (1997) The *glossy1* locus of maize and an epidermis-specific cDNA from *Kleinia odora* define a class of receptor-like proteins required for the normal accumulation of cuticular waxes. Plant Physiol 113: 1091–1100

Hayes HK, Brewbaker HE (1928) *Glossy* seedlings in maize. Am Nat 62: 228–235

Hose E, Clarkson DT, Steudle E, Schreiber L, Hartung W (2001) The exodermis: A variable apoplastic barrier. J Exp Bot 52: 2245–2264

Huang J, Xue C, Wang H, Wang L, Schmidt W, Shen R, Lan P (2017) Genes of ACYL CARRIER PROTEIN family show different expression profiles and overexpression of ACYL CARRIER PROTEIN 5 modulates fatty acid composition and enhances salt stress tolerance in Arabidopsis. Front Plant Sci 8:987

Jiao Y, Peluso P, Shi J, Liang T, Stitzer MC, Wang B, Campbell MS, Stein JC, Wei X, Chin C-S, et al (2017) Improved maize reference genome with single-molecule technologies. Nature 546: 524–527

Jun JH, Song Z, Liu Z, Nikolau BJ, Yeung ES, Lee YJ (2010) High-spatial and high-mass resolution imaging of surface metabolites of *Arabidopsis thaliana* by laser desorption-ionization mass spectrometry using colloidal silver. Anal Chem 82: 3255–3265

Kim D, Langmead B, Salzberg SL (2015) HISAT: A fast spliced aligner with low memory requirements. Nat Methods 12: 357–360

Kim H, Lee SB, Kim HJ, Min MK, Hwang I, Suh MC (2012) Characterization of Glycosylphosphatidylinositol-Anchored Lipid Transfer Protein 2 (LTPG2) and overlapping function between LTPG/LTPG1 and LTPG2 in cuticular wax export or accumulation in *Arabidopsis thaliana*. Plant Cell Physiol 53: 1391–1403

Kim J-JP, Miura R (2004) Acyl-CoA dehydrogenases and acyl-CoA oxidases. Structural basis for mechanistic similarities and differences. Eur J Biochem 271: 483–493

Kim JY, Song JT, Seo HS (2017) COP1 regulates plant growth and development in response to light at the post-translational level. J Exp Bot 68: 4737–4748

Kim RJ, Kim HJ, Shim D, Suh MC (2016) Molecular and biochemical characterizations of the monoacylglycerol lipase gene family of *Arabidopsis thaliana*. Plant J 85: 758–771

Kobayashi K, Masuda T (2016) Transcriptional regulation of tetrapyrrole biosynthesis in *Arabidopsis thaliana*. Front Plant Sci 7: 1811

Kolattukudy PE, Walton TJ (1973) The biochemistry of plant cuticular lipids. Prog Chem Fats Other Lipids 13: 119–175

Krijthe JH (2015) Rtsne: T-Distributed Stochastic Neighbor Embedding using a Barnes-Hut implementation. R package version 0.16, https://github.com/jkrijthe/Rtsne

Kunst L, Samuels L (2009) Plant cuticles shine: Advances in wax biosynthesis and export. Curr Opin Plant Biol 12: 721–727

Langfelder P, Horvath S (2008) WGCNA: An R package for weighted correlation network analysis. BMC Bioinformatics 9: 559

Lauter N, Kampani A, Carlson S, Goebel M, Moose SP (2005) microRNA172 down-regulates *glossy15* to promote vegetative phase change in maize. Proc Natl Acad Sci USA 102: 9412–9417

Legris M, Ince YÇ, Fankhauser C (2019) Molecular mechanisms underlying phytochrome-controlled morphogenesis in plants. Nat Commun 10: 5219

Li F-S, Phyo P, Jacobowitz J, Hong M, Weng J-K (2019a) The molecular structure of plant sporopollenin. Nat Plants 5: 41–46

Li L, Du Y, He C, Dietrich CR, Li J, Ma X, Wang R, Liu Q, Liu S, Wang G, et al (2019b) Maize *glossy6* is involved in cuticular wax deposition and drought tolerance. J Exp Bot 70: 3089–3099

Li L, Li D, Liu S, Ma X, Dietrich CR, Hu H-C, Zhang G, Liu Z, Zheng J, Wang G, et al (2013) The Maize *glossy13* gene, cloned via BSR-Seq and Seq-Walking encodes a putative ABC transporter required for the normal accumulation of epicuticular waxes. PLoS One 8: e82333

Li Y, Beisson F, Koo AJK, Molina I, Pollard M, Ohlrogge J (2007) Identification of acyltransferases required for cutin biosynthesis and production of cutin with suberin-like monomers. Proc Natl Acad Sci USA 104: 18339–18344

Liaw A, Wiener M (2002) Classification and regression by randomForest. R News 2: 18–22

Li-Beisson Y, Shorrosh B, Beisson F, Andersson MX, Arondel V, Bates PD, Baud S, Bird D, DeBono A, Durrett TP, et al (2010) Acyl-lipid metabolism. Arabidopsis Book 8: e0133

Lin M, Matschi S, Vasquez M, Chamness J, Kaczmar N, Baseggio M, Miller M, Stewart EL, Qiao P, Scanlon MJ, et al (2020) Genome-wide association study for maize leaf cuticular conductance identifies candidate genes involved in the regulation of cuticle development. G3 10: 1671–1683

Lin M, Qiao P, Matschi S, Vasquez M, Ramstein GP, Bourgault R, Mohammadi M, Scanlon MJ, Molina I, Smith LG, et al (2022) Integrating GWAS and TWAS to elucidate the genetic architecture of maize leaf cuticular conductance. Plant Physiol 189: 2144–2158

Liu S, Dietrich CR, Schnable PS (2009) DLA-based strategies for cloning insertion mutants: Cloning the *gl4* locus of maize using *Mu* transposon tagged alleles. Genetics 183: 1215–1225

Liu X, Bourgault R, Galli M, Strable J, Chen Z, Feng F, Dong J, Molina I, Gallavotti A (2021) The FUSED LEAVES1-*ADHERENT1* regulatory module is required for maize cuticle development and organ separation. New Phytol 229: 388–402

Liu Y, Gu H-Y, Zhu J, Niu Y-M, Zhang C, Guo G-L (2019) Identification of hub genes and key pathways associated with bipolar disorder based on weighted gene co-expression network analysis. Front Physiol 10: 1081

Loneman DM, Peddicord L, Al-Rashid A, Nikolau BJ, Lauter N, Yandeau-Nelson MD (2017) A robust and efficient method for the extraction of plant extracellular surface lipids as applied to the analysis of silks and seedling leaves of maize. PLoS One 12: e0180850

Lu Y-H, Arnaud D, Belcram H, Falentin C, Rouault P, Piel N, Lucas M-O, Just J, Renard M, Delourme R, et al (2013) A dominant point mutation in a RINGv E3 ubiquitin ligase homoeologous gene leads to cleistogamy in *Brassica napus*. Plant Cell 24: 4875–4891

McFarlane HE, Watanabe Y, Yang W, Huang Y, Ohlrogge J, Samuels AL (2014) Golgi- and trans-golgi network-mediated vesicle trafficking is required for wax secretion from epidermal cells. Plant Physiol 164: 1250–1260

Moose SP, Sisco PH (1994) *Glossy15* controls the epidermal juvenile-to-adult phase transition in maize. Plant Cell 16: 1343–1355

Perera MADN, Qin W, Yandeau-Nelson M, Fan L, Dixon P, Nikolau BJ (2010) Biological origins of normal-chain hydrocarbons: A pathway model based on cuticular wax analyses of maize silks. Plant J 64: 618–632

Petit J, Bres C, Reynoud N, Lahaye M, Marion D, Bakan B, Rothan C (2021) Unraveling cuticle formation, structure, and properties by using tomato genetic diversity. Front Plant Sci 12: 778131

Post-Beittenmiller D (1996) Biochemistry and molecular biology of wax production in plants. Annu Rev Plant Physiol Plant Mol Biol 47: 405–430

Qiao P, Bourgault R, Mohammadi M, Matschi S, Philippe G, Smith LG, Gore MA, Molina I, Scanlon MJ (2020) Transcriptomic network analyses shed light on the regulation of cuticle development in maize leaves. Proc Natl Acad Sci USA 117: 12464–12471

R Core Team (2019) R: A language and environment for statistical computing. R Foundation for Statistical Computing.

Reynoud N, Geneix N, Petit J, D’Orlando A, Fanuel M, Marion D, Rothan C, Lahaye M, Bakan B (2022) The cutin polymer matrix undergoes a fine architectural tuning from early tomato fruit development to ripening. Plant Physiol 190: 1821–1840

Reynoud N, Petit J, Bres C, Lahaye M, Rothan C, Marion D, Bakan B (2021) The complex architecture of plant cuticles and its relation to multiple biological functions. Front Plant Sci 10: 782773

Riederer M, Schreiber L (2001) Protecting against water loss: Analysis of the barrier properties of plant cuticles. J Exp Bot 52: 2023–2032

la Rocca N, Manzotti PS, Cavaiuolo M, Barbante A, Dalla Vecchia F, Gabotti D, Gendrot G, Horner DS, Krstajic J, Persico M, et al (2015) The maize *fused leaves1* (*fdl1*) gene controls organ separation in the embryo and seedling shoot and promotes coleoptile opening. J Exp Bot 66: 5753–5767

Rocha J, Nitenberg M, Girard-Egrot A, Jouhet J, Maréchal E, Block MA, Breton C (2018) Do galactolipid synthases play a key role in the biogenesis of chloroplast membranes of higher plants? Front Plant Sci 9: 126

Schnable PS, Stinard P, Wen T-J, Heinen S, Weber DF, Schneerman M, Zhang L, Hansen JD, Nikolau BJ (1994) The genetics of cuticular wax biosynthesis. Maydica 39: 279–287

Schwacke R, Ponce-Soto GY, Krause K, Bolger AM, Arsova B, Hallab A, Gruden K, Stitt M, Bolger ME, Usadel B (2019) MapMan4: a refined protein classification and annotation framework applicable to multi-omics data analysis. Mol Plant 12: 879–892

Seale M (2020) The fat of the land: Cuticle formation in terrestrial plants. Plant Physiol 184: 1622–1624

Shannon P, Markiel A, Ozier O, Baliga NS, Wang JT, Ramage D, Amin N, Schwikowski B, Ideker T (2003) Cytoscape: A software environment for integrated models of biomolecular interaction networks. Genome Res 13: 2498–2504

Shepherd T, Wynne Griffiths D (2006) The effects of stress on plant cuticular waxes. New Phytol 171: 469–499

Song L, Langfelder P, Horvath S (2013) Random generalized linear model: A highly accurate and interpretable ensemble predictor. BMC Bioinformatics 14: 5

Stein SE (1999) An integrated method for spectrum extraction and compound identification from gas chromatography/mass spectrometry data. J Am Soc Mass Spectrom 10: 770–781

Sturaro M, Hartings H, Schmelzer E, Velasco R, Salamini F, Motto M (2005) Cloning and characterization of *GLOSSY1*, a maize gene involved in cuticle membrane and wax production. Plant Physiol 138: 478–489

Sun S, Zhou Y, Chen J, Shi J, Zhao H, Zhao H, Song W, Zhang M, Cui Y, Dong X, et al (2018) Extensive intraspecific gene order and gene structural variations between Mo17 and other maize genomes. Nat Genet 50: 1289–1295

Supek F, Bošnjak M, Škunca N, Šmuc T (2011) REVIGO Summarizes and visualizes long lists of gene ontology terms. PLoS One 6: e21800

Suresh K, Zeisler-Diehl V v., Wojciechowski T, Schreiber L (2022) Comparing anatomy, chemical composition, and water permeability of suberized organs in five plant species: wax makes the difference. Planta 256: 60

Teramoto H, Ono T, Minagawa J (2001) Identification of *Lhcb* gene family encoding the light-harvesting chlorophyll-a/b proteins of photosystem II in *Chlamydomonas reinhardtii*. Plant Cell Physiol 42: 849–856

Vierstra RD (2009) The ubiquitin–26S proteasome system at the nexus of plant biology. Nat Rev Mol Cell Biol 10: 385–397

Wan H, Liu H, Zhang J, Lyu Y, Li Z, He Y, Zhang X, Deng X, Brotman Y, Fernie AR, et al (2020) Lipidomic and transcriptomic analysis reveals reallocation of carbon flux from cuticular wax into plastid membrane lipids in a *glossy* “Newhall” navel orange mutant. Hortic Res 7: 41

WANG X, DEVAIAH S, ZHANG W, WELTI R (2006) Signaling functions of phosphatidic acid. Prog Lipid Res 45: 250–278

Wang Z, Tian X, Zhao Q, Liu Z, Li X, Ren Y, Tang J, Fang J, Xu Q, Bu Q (2018) The E3 ligase DROUGHT HYPERSENSITIVE negatively regulates cuticular wax biosynthesis by promoting the degradation of transcription factor ROC4 in rice. Plant Cell 30: 228–244

Wimalanathan K, Friedberg I, Andorf CM, Lawrence-Dill CJ (2018) Maize GO annotation—Methods, evaluation, and review (maize-GAMER). Plant Direct 2: e00052

Xia Y, Gao Q-M, Yu K, Lapchyk L, Navarre D, Hildebrand D, Kachroo A, Kachroo P (2009) An intact cuticle in distal tissues is essential for the induction of systemic acquired resistance in plants. Cell Host Microbe 5: 151–165

Yang B, Liu J, Ma X, Guo B, Liu B, Wu T, Jiang Y, Chen F (2017) Genetic engineering of the Calvin cycle toward enhanced photosynthetic CO2 fixation in microalgae. Biotechnol Biofuels 10: 229

Yang Y, Shi J, Chen L, Xiao W, Yu J (2022) ZmEREB46, a maize ortholog of Arabidopsis WAX INDUCER1/SHINE1, is involved in the biosynthesis of leaf epicuticular very-long-chain waxes and drought tolerance. Plant Science 321: 111256

Yeats TH, Rose JKC (2013) the formation and function of plant cuticles. Plant Physiol 163: 5–20

Yeh S-Y, Lin H-H, Chang Y-M, Chang Y-L, Chang C-K, Huang Y-C, Ho Y-W, Lin C-Y, Zheng J-Z, Jane W-N, et al (2021) Maize Golden2-like transcription factors boost rice chloroplast development, photosynthesis, and grain yield. Plant Physiol 188: 442–459

Zaman M, Kleineidam K, Bakken L, Berendt J, Bracken C, Butterbach-Bahl K, Cai Z, Chang SX, Clough T, Dawar K, et al (2021) Greenhouse gases from agriculture. Measuring emission of agricultural greenhouse gases and developing mitigation options using nuclear and related techniques. Springer International Publishing, Cham, pp 1–10

Zhao X, Wei J, He L, Zhang Y, Zhao Y, Xu X, Wei Y, Ge S, Ding D, Liu M, et al (2019) Identification of fatty acid desaturases in maize and their differential responses to low and high temperature. Genes 10: 445

Zheng J, He C, Qin Y, Lin G, Park WD, Sun M, Li J, Lu X, Zhang C, Yeh C, et al (2018) Co-expression analysis aids in the identification of genes in the cuticular wax pathway in maize. Plant J 97: 530–542

